# Simple Sequence Repeats Mediate Phase Variation of the Mucoid Phenotype in Hypervirulent *Klebsiella pneumoniae*

**DOI:** 10.1101/2025.09.12.675794

**Authors:** Oliwia Stawarska, Aarabhi Ravikrishnan Nair, Ania W. Y. Man, Luke R. Green, Ozcan Gazioglu, Francesco Flandi, Margaret M. C. Lam, Kathryn E. Holt, Hasan Yesilkaya, Marco R. Oggioni, Jose A. Bengoechea, Charlotte Odendall, Alex J. McCarthy, Joseph J. Wanford

## Abstract

Hypervirulent *Klebsiella pneumoniae* (HV*Kp*) express a thick mucoid capsular polysaccharide which is essential for virulence and survival in the bloodstream but inhibits epithelial adhesion and invasion. The molecular mechanisms regulating capsule expression to rapidly adapt to these distinct niches are unknown. Here, we show that high frequency mutations in a poly-G repeat in the regulator of mucoid phenotype gene, *rmpA*, mediates reversible ON/OFF phase variation of the mucoid phenotype. We find that these mutations generate subpopulations of bacteria with reduced capsule expression and mucoidy. *In vitro* and *in vivo* infection models reveal that phase variation to the OFF state facilitates epithelial cell infection while ON switching enhances resistance to the bactericidal activity of human serum and systemic spread to the liver and spleen in a murine infection model. Genomic analysis reveals that the phase variation mechanism is a conserved feature of all HV*Kp* and that additional novel DNA repeats in *rmpD* and *rmpC* drive different combinations of capsule mucoidy and hyperexpression phenotypes. These findings establish a mechanistic basis for how HV*Kp* occupy such a diverse array of niches during infection, with important implications for the design and implementation of *Kp*-specific vaccines and therapies.

## Introduction

The Gram-negative bacterium *Klebsiella pneumoniae* (*Kp*) is a human commensal, and pathogen exerting a major burden on world health^1^. The hypervirulent pathotype (HVKp) is of critical concern as it causes disease in immunocompetent individuals with high mortality rates ^2,3^. This pathotype asymptomatically colonises the human gastrointestinal tract but can disseminate systemically to cause severe bloodstream infections and pyogenic tissue abscesses ^4–6^. Since its first description in Taiwan, HV*Kp* has undergone epidemic spread throughout Asia and is disseminating across the globe^7–9^. The emergence of phenotypically convergent strains exhibiting markers of both hypervirulence and multi-drug resistance are a major cause for concern ^10,11^. Development of new therapeutic strategies and preventative vaccines to combat this threat requires an in depth understanding of mechanisms of virulence.

HV*Kp* are characterised by expression of a thick extracellular polysaccharide capsule which is required for virulence in both systemic and pulmonary infection models ^12–17^. Capsule promotes resistance to killing by the serum complement system and antimicrobial peptides, but contrastingly forms a barrier to adhesion to and invasion of phagocytic and non-phagocytic cells ^18–25^. Indeed, despite being classically defined as an extracellular pathogen, a growing body of evidence demonstrates that HV*Kp* can survive intracellularly to facilitate immune evasion and recalcitrance to antibiotic treatment ^26–29^. The capacity of these highly encapsulated organisms to both colonise the epithelium, survive in the bloodstream, and to occupy both the intra– and extra-cellular space is an unaddressed paradox, suggestive that unknown mechanisms regulate capsule expression and mucoidy during HV*Kp* infection.

HV*Kp* predominantly express capsules of serotype K1 and K2 which are both highly expressed and display long polysaccharide chain lengths conferring a physically mucoid phenotype ^30^. These two phenotypes are genetically separable and are conferred by the *rmp* locus, consisting of *rmpA, rmpD,* and *rmpC.* This locus is primarily encoded by two virulence plasmids – termed KpVP-1 and KpVP-2 – in addition to integrative conjugative elements (ICEs) inserted into the chromosome ^31–34^. RmpA is a transcription factor and autoregulator of the operon ^35,36^. RmpD interacts with the capsule biosynthesis component Wzc to modulate capsule polymer length and confers the mucoid phenotype ^37^. RmpC is a transcription factor which positively regulates the capsule biosynthesis locus driving hyper-capsule expression ^38^. Capsule expression and chain length are further modulated by a network of additional transcriptional regulators which respond to environmental cues including temperature, pH and nutrient availability ^39–42^. However, the specific mechanisms that promote rapid switching of hypermucoid capsule expression during HV*Kp* infection remain undefined.

Phase variation involves stochastic, high frequency, reversible, ON/OFF switches in gene expression with consequent generation of phenotypic heterogeneity in otherwise clonal populations ^43,44^. High frequency phase variation enables bacterial pathogens to maintain fitness by overcoming the restrictive effects of selective and non-selective bottlenecks during transmission between and within host compartments ^44–48^. By pre-emptively generating large numbers of variants during replication, pathogens can ‘hedge their bets’ to adapt to fluctuating host and environmental selection pressures. The best characterised mechanism of phase variation is exemplified by intragenic simple sequence repeats (SSRs) in *Neisseria* and *Campylobacter ssp* ^49–52^, in which repetitive DNA stretches are prone to DNA polymerase slippage and localised hypermutation during replication, leading to high frequency insertion/deletion (InDel) mutations. These mutations often produce frameshifts and result in switching SSR-containing genes into an OFF state ^53,54^. Subsequent rounds of slippage and hypermutation during replication allow for regain of gene function and reversible switches between pathogenic phenotypes. Whether phase variation of the mucoid phenotype drives adaptation of HV*Kp* to different within-host niches remains unexplored.

In this study, we demonstrate that a poly-G repeat in the *rmpA* gene mediates high-frequency ON/OFF phase variation of the mucoid phenotype during host cell infection. We show that this mechanism is conserved across all HV*Kp*, and that phase variation mediates a phenotypic switch between high and low epithelial infection states with consequent impacts on systemic virulence. These results support a model whereby the rapid generation of phenotypic diversity through mutations at repetitive DNA prime *Kp* populations to adapt to fluctuating selection for capsule expression. Our study provides critical insights into how this major pathogen remodels expression of its capsule during infection, with implications for therapy and rationale vaccine design.

## Results

### HV*Kp* host cell infection is associated with loss of the mucoid phenotype

The expression of HV*Kp* capsule is reported to have an inhibitory effect on HV*Kp* adhesion to host cells ^24,55,56^. This led us to hypothesise that a clonal population of HV*Kp* contains a subpopulation expressing reduced mucoid capsule that facilitates their initial adhesion to and invasion of host cells. To evaluate this, we cultured the serotype K1 Clonal Group (CG) 23 strain SGH10 (Table 1), infected A549 lung epithelial cells and THP-1 macrophage-like cells (both cell types of which are encountered by HV*Kp* during infection), and quantified the mucoid phenotype of the adherent and invasive populations using a high throughput sedimentation resistance assay (Figure 1a; Figure S1). We observed a large population bottleneck during both bacterial adhesion and internalisation, with only ∼0.0001% of the inoculum being internalised by A549 cells, and ∼0.01% of bacteria phagocytosed by THP-1 cells (Figure 1b). Quantification of the mucoid phenotype of multiple individual colonies from the input, adherent and invasive populations revealed that SGH10 populations underwent a significant loss of mucoidy during infection (Figure 1c). Given the genetic diversity of *Kp,* we next studied an additional HV*Kp* strain, SB4538 with capsule serotype K2 from CG25 (Table 1). Compared with SGH10, we observed a similar population bottleneck during adhesion and invasion of both cell lines with SB4538 (Figure 1d). Strikingly however, strain SB4538 underwent a dramatic and stepwise loss of mucoidy during both adhesion and invasion (Figure 1e). Given the strength of selection for loss of mucoidy observed in strain SB4538, we re-cultured 8 cell-associated variants from this background and subjected them to a second round of infection. This indicated that 5 of 8 strains retained an enhanced capacity to adhere to A549 cells compared with the wildtype, and all 8 strains retained an enhanced invasion phenotype (Figure 1f-g). Together, these data indicate that HV*Kp* populations exhibit heterogeneity in their capacity to adhere to and invade host cells.

**Figure 1.**
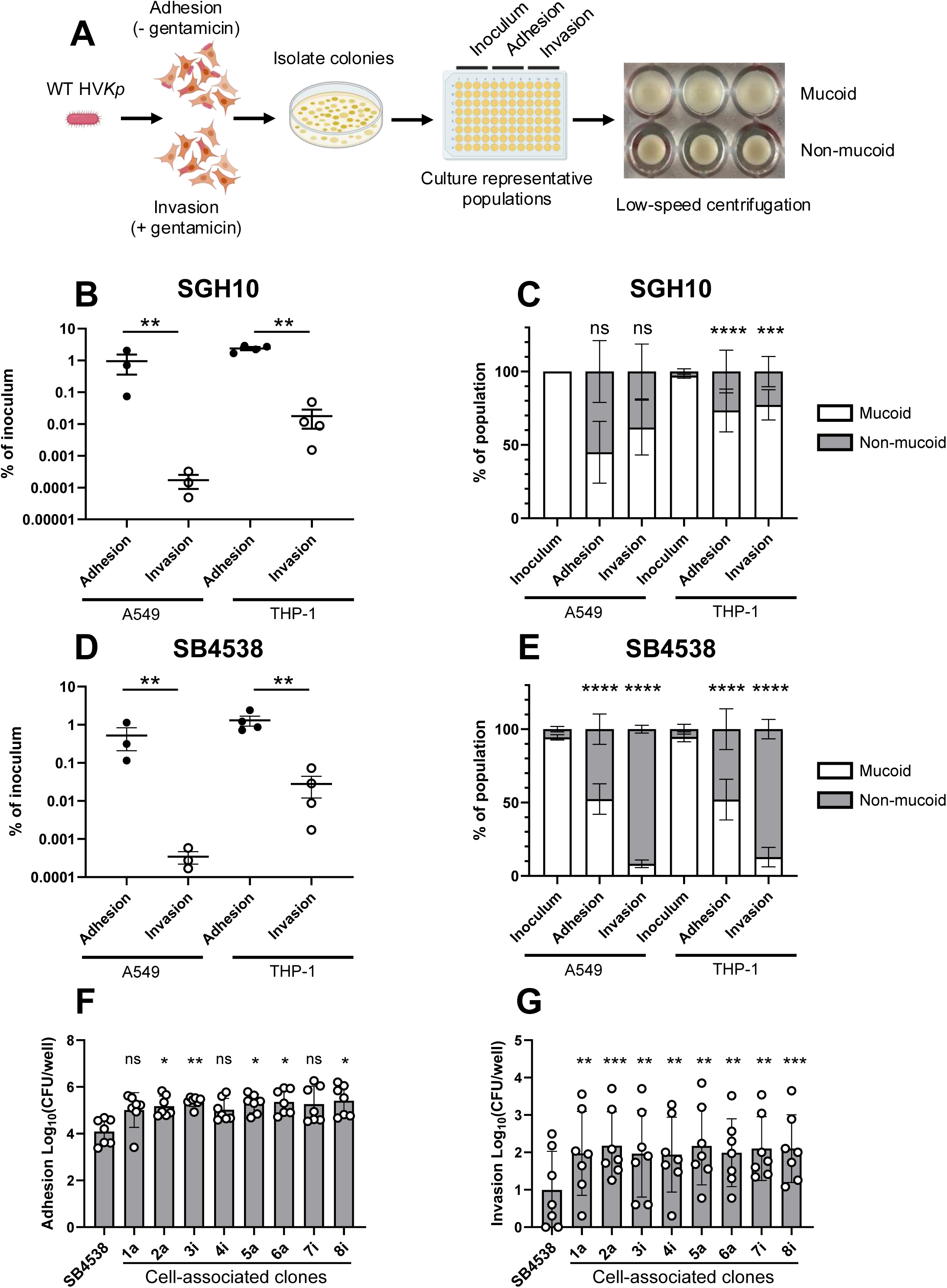
HV*Kp* host cell infection is associated with loss of the mucoid phenotype. (A) Schematic of the experimental workflow to measure population scale changes in mucoidy during infection. (B) Burdens of K1 strain SGH10 isolated following adhesion to and invasion of A549 and THP-1 cells. (C) Percentage of SGH10 colonies displaying the mucoid phenotype as measured by a high throughput sedimentation resistance assay following isolation of adhesive and invasive bacteria from A549 and THP-1 cells. (D) Burdens of K2 strain SB4538 isolated following adhesion to and invasion of A549 and THP-1 cells. (E) Percentage of SB4538 colonies displaying the mucoid phenotype as measured by a high throughput sedimentation resistance assay following isolation of adhesive and invasive bacteria from A549 and THP-1 cells. (F) Bacterial burdens isolated from adherent and invasive fractions of A549 epithelial cells infected with wild type SB4538 and non-mucoid colonies that were enriched from a first round of cell infections. Data are expressed as the mean total CFU per well from at least 3 independent biological replicates. Statistical significance was determined in panels B, C, and D using a one-way ANOVA, and in panels C and E using a Generalised Linear Mixed Model (see materials and methods). ****; P> 0.00005, ***; P>0.0005, **; P>0.005, *; P>0.05, ns; not significant.

**Table 1.** Bacterial strains and plasmids used in this study.

| Strain / plasmid | Description | *rmpA* genotype | Source |
| --- | --- | --- | --- |
| *K. pneumoniae* |  |  |  |
| SGH10 | Serotype K1, ST23 | WT, functional (*rmpA* G10) | <sup>7</sup> |
| SB4538 | Serotype K2, ST25 | WT, functional (*rmpA* G10) | Sylvain Brisse |
| Kp43816 | Serotype K2 type strain. | WT, functional (*rmpA* G10) | Jose Bengoechea |
| Kp43816 Δ*manC* Δ*glf* | Capsule and O-antigen double mutant derivative of Kp43816 | WT, functional (*rmpA* G10) | Jose Bengoechea |
| Plasmids |  |  |  |
| pRmpA-kan | *rmpA* allele 2 subcloned into Puc57-Kan | WT, functional (*rmpA* G10) | This study |
| pBR322 | AmpR, TetR cloning vector. | N/A | Addgene |
| pBR322-*rmp*-111 | *rmp1* and 500bp upstream from SGH10 cloned into AatII and PstI restriction sites of pBR322 | See description. | This study |
| pBR322-*rmp*-011 | As above, but with *rmpA* G9 repeat | See description. | This study |
| pBR322-*rmp*-101 | As above but with *rmpD* T8 repeat | See description. | This study |
| pBR322-*rmp*-110 | As above but with *rmpC* G8 repeat | See description. | This study |

WT; wildtype, G10; poly-G repeat consisting of 10 monomeric units. ST; sequence type.

### Infection selects for inactivating mutations in *rmpA* and loss of the large virulence plasmid

As our adhesion and invasion data indicated that host cell infection selects for loss of mucoidy, we hypothesised that mutations in regulatory genes of the capsule operon were driving switching from a mucoid to non-mucoid phenotype during epithelial cell infection. To test this, five cell-associated clones from each strain background were subjected to hybrid whole genome sequencing with Illumina and Oxford Nanopore Technology (ONT) to generate hybrid assemblies. Analysis of virulence gene content with Kleborate and comparison to the reference genome indicated two candidate genetic routes to loss of mucoidy during epithelial cell infection (Figure 2a; Table 2). Genome sequences obtained from two clones in both strain background assembled into a single contig. These assemblies were missing an additional 200 kbp contig found in the WT sequence and lacked plasmid-associated virulence genes and the KpVP-1 plasmid replicon, indicating that these variants had lost the large virulence plasmid. Contrastingly, the remaining 3/5 mutants in each strain background harboured an insertion mutation at a poly-G repeat in the *rmpA* gene (Figure 2b).

**Figure 2.**
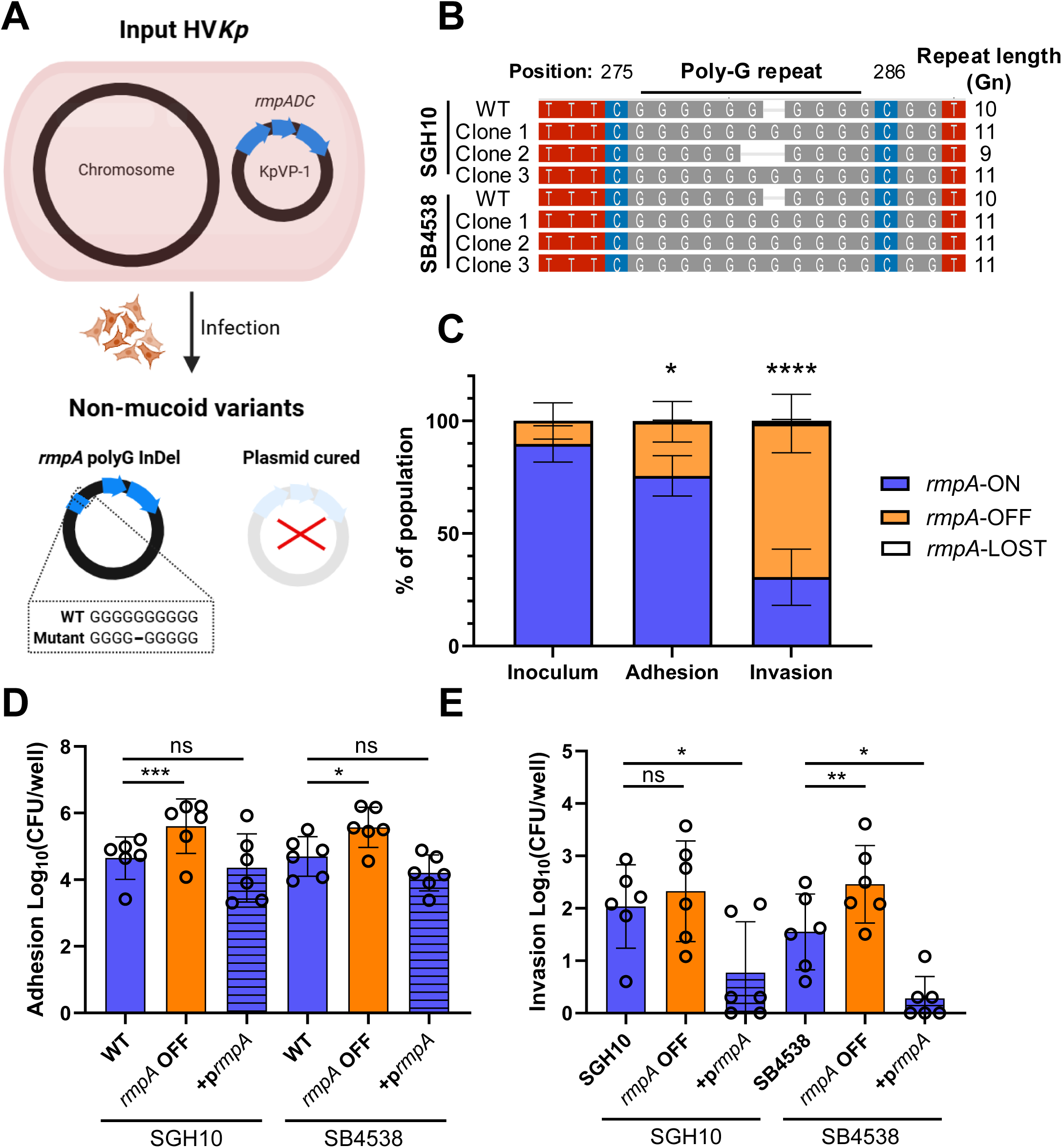
Loss of mucoidy is driven by *rmpA* truncation and loss of the large virulence plasmid. (A) Schematic depicting the two candidate routes to loss of mucoidy during epithelial cell infection. (B) Alignment of the *rmpA* sequence from WT SGH10, SB4538, and 3 invasive clones in each strain background indicating mutations at the *rmpA* poly-G repeat. (C) PCR-based analysis of *rmpA* truncation and *rmpA* loss in the inoculum and adhesive and invasive fractions isolated from A549 infected with strain SB4538 (see methods). The fraction of *rmpA*-ON colonies is indicated by blue bars, *rmpA-*OFF colonies by orange bars, and *rmpA*-LOST colonies by white bars. Data are representative of 3 independent experiments, and statistical significance was determined by comparing the proportion of colonies with each genotype using a Generalise Linear Mixed Model. ***; P>0.0005, **; P>0.005, *; P>0.05, ns; not significant. (D) Adhesive and (E) invasive bacterial burdens from A549 cells following infection with WT HV*Kp*, their poly-G mutated *rmpA*-OFF variants, and strains complemented with a functional copy of *rmpA.* Data are representative of 6 independent experiments and statistical significance were determined using an ordinary 1-way ANOVA. ns; non-significant, *; P>0.05, **; P>0.005, ***; P>0.0005.

**Table 2.** Genomic characteristics of non-mucoid HV*Kp* clones.

| Strain | ST | Inc Type | Aerobactin | Salmoachelin | *rmpA* | Poly-G Length |
| --- | --- | --- | --- | --- | --- | --- |
| SGH10 WT | 23 | repB_KLEB | + (*iuc* 1) | + (*iro* 1) | + (2) | 10 |
| Clone 1 | 23 | - | - | - | - | N/A |
| Clone 2 | 23 | - | - | - | - | N/A |
| Clone 3 | 23 | repB_KLEB | + (*iuc* 1) | + (*iro* 1) | + (4), Truncated (47%) | 11 |
| Clone 4 | 23 | repB_KLEB | + (*iuc* 1) | + (*iro* 1) | + (5), Truncated (54%) | 9 |
| Clone 5 | 23 | repB_KLEB | + (*iuc* 1) | + (*iuc* 1) | + (4), Truncated (47%) | 11 |
| SB4538 WT | 25 | IncHI1B | + (*iuc* 1) | + (*iro* 1) | + (2) | 10 |
| Clone 1 | 25 | - | - | - | - | N/A |
| Clone 2 | 25 | - | - | - | - | N/A |
| Clone 3 | 25 | IncHI1B(pNDM-MAR) repB_KLEB | + (*iuc* 1) | + (*iro* 1) | + (4), Truncated (47%) | 11 |
| Clone 4 | 25 | IncHI1B(pNDM-MAR) repB_KLEB | + (*iuc* 1) | + (*iro* 1) | + (4), Truncated (47%) | 11 |
| Clone 5 | 25 | IncHI1B(pNDM-MAR) repB_KLEB | + (*iuc* 1) | + (*iro* 1) | + (4), Truncated (47%) | 11 |

Alleles numbers and the percentage of the gene functionally expressed as defined by Kleborate are shown in brackets.
+, gene present, -, gene absent. N/A; not applicable. ST; sequence type.

Given that repetitive DNA sequences are prone to localised hypermutation and accumulation of insertion/deletion mutations (InDels) at several orders of magnitude higher than global mutation rates ^57^, we hypothesised that *rmpA* mutations would be highly frequent and the major mutation present in cell-associated *Kp.* To measure *rmpA* mutations and loss of the virulence plasmid at the population level, we developed a PCR-based assay, involving amplification of the repeat region in *rmpA* using fluorescent primers on DNA derived from multiple individual colony lysates (Figure S2). We included *rmpC* (additional virulence plasmid control) and *wzi* (a chromosomal DNA control) in a multiplex reaction to infer plasmid loss and template DNA quality, respectively. We excluded the *rmpA* homolog, *rmpA2*, because although it has a poly-G repeat, it typically contains stop codons downstream that consistently truncate the gene, regardless of poly-G repeat length (Fig S3). Based on multiplex PCR results, colonies were subsequently coded as *rmpA*-ON (WT length), *rmpA*-OFF (altered amplicon length due to repeat expansion/contraction) or *rmpA*-LOST (*wzi* positive and *rmpAC* negative). Applying this assay to colonies obtained from replicate infections of A549 cells with SB4538, where we saw the clearest reduction in mucoidy, we observed a stepwise increase in the number of *rmpA*-OFF colonies isolated from adhesive and invasive populations, respectively, with very few *rmpA*-LOST colonies recovered in these assays (Figure 2c). As our data reported that HV*Kp* with *rmpA*-OFF mutations display increased efficiency in cellular adhesion and invasion, it would be expected that a strain with constitute *rmpA* expression would be less efficient at host cell invasion. Indeed, complementation of *rmpA*-OFF mutants in both strain backgrounds with a functional copy of *rmpA* on a high copy number plasmid reduced adhesion to A549 cells to levels comparable to the WT (Figure 2d), and significantly reduced levels of intracellular invasion (Figure 2e). Together, these data indicate that HV*Kp* host cell infection is associated with inactivating mutations in an intragenic poly-G tract within *rmpA*, and occasional loss of the large virulence plasmid.

### *rmpA* poly-G repeat mutations drive population variation in mucoid capsule expression during non-selective growth

To determine the functional consequences of *rmpA* inactivation, we compared the predicted structure of RmpA from WT HV*Kp* and *in cellulo-*isolated RmpA-truncation mutants. This revealed that the poly-G repeat encodes amino acids localised to a flexible linker between two distinct domains of RmpA (Figure 3a), and that truncation led to loss of the region containing the putative LuxR-like DNA-binding domain. This indicated that these poly-G tract mutations likely lead to loss of *rmpA* function.

**Figure 3.**
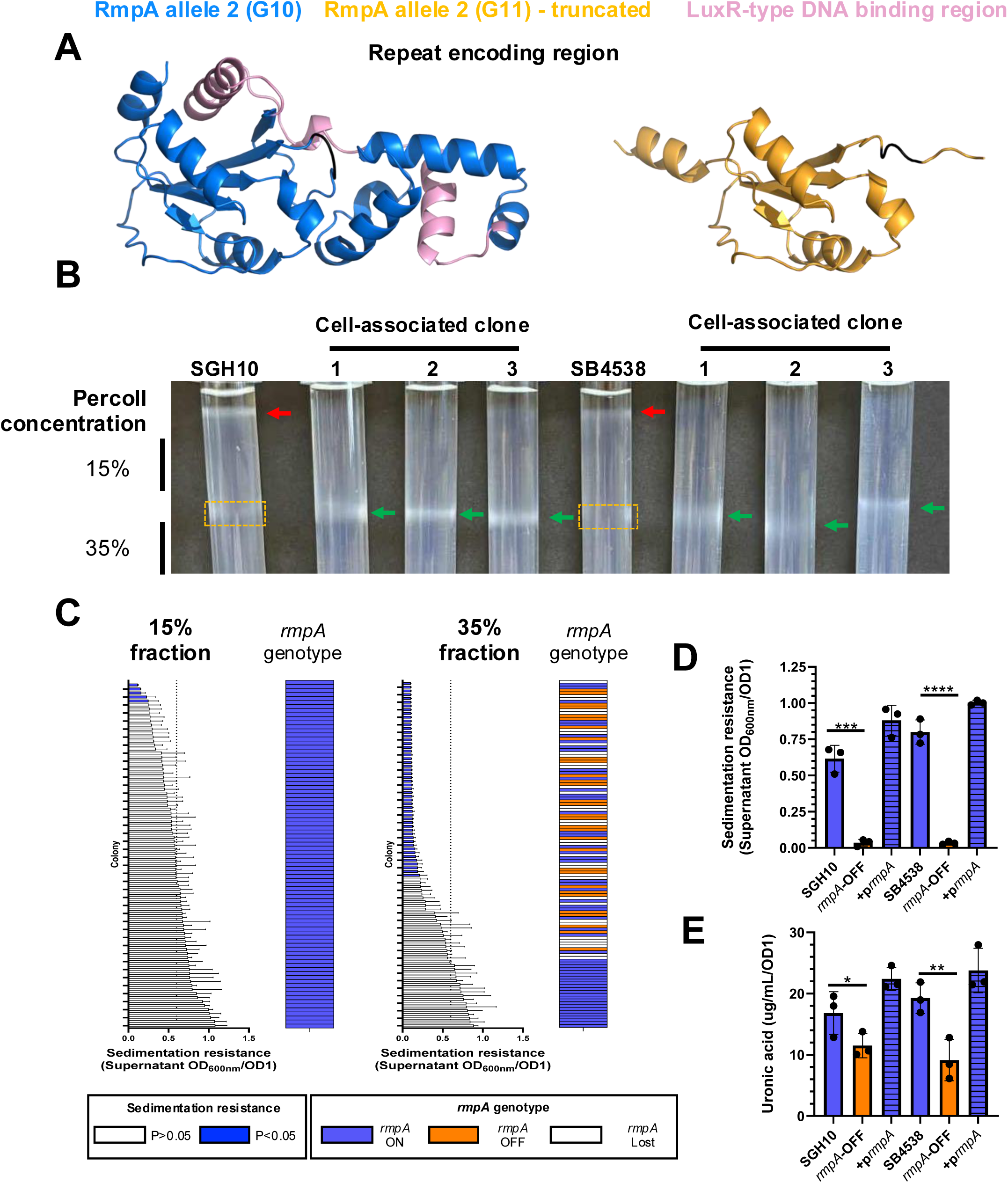
Mutations in the *rmpA* poly-G repeat generate population-level variation in capsule expression. (A) Alphafold model of RmpA protein produced by *rmpA* allele 2 (blue), and the G11 truncated variant isolates from epithelial cells (orange). The LuxR-type DNA binding region (amino acids 110-156) is shown in pink, and the region encoded by the poly-G repeat is shown in black. (B) Localisation of WT HV*Kp* and invasive clones isolated from A549 epithelial cells in discontinuous Percoll gradients. WT fractions are indicated by red arrows; mutant fractions are indicated by green arrows. High– and low-density fractions in WT HV*Kp* gradients used to isolate colonies for downstream analysis are indicated by yellow dashed boxes. (C) Correlation of sedimentation resistance (bar graph), with the presence and expression state of *rmpA* derived from PCR fragment size analysis of individual colonies (heatmap; see materials and methods). Each bar and row of the heatmap corresponds to an individual colony isolated from the top (left) or bottom (right) Percoll fractions. The dotted line indicates the plate average, and bars coloured in blue are significantly different to the WT mucoid control. Orange blocks indicate functional expression of the gene, blue blocks indicate a frameshift mutation, and white blocks indicate absence of the gene. (D) Sedimentation resistance assay of SGH10 and SB4538 WT strains (blue bars), *rmpA* OFF phase variants (orange bars), and OFF variants transformed with a functional *rmpA* complementation plasmid (blue thatched bars) (n=3). (E) Measurement of capsule uronic acid in strains as in D. Error bars indicate the standard deviation. Statistical significances were determined by ordinary one-way ANOVA with Dunnett’s post-hoc test in all cases with comparison to the WT strain. ****; P>0.00005, ***; P>0.0005, **; P>0.005, *; P>0.05.

Given that *rmpA* is required for both capsule mucoidy and hyperexpression through regulation of *rmpD* and *rmpC,* respectively^37,38^, we hypothesised that these mutations would drive population level variation in capsule expression. To test this, we separated saturated overnight cultures of WT HV*Kp* and *in cellulo*-isolated *rmpA* mutants, predicted to have differential capsule expression, using a discontinuous Percoll gradient (Figure 3b)^41^. This revealed that WT HV*Kp* localised to the top of the gradient, whereas *in cellulo*-isolated *rmpA* mutants in both strain backgrounds displayed an intermediate capsule phenotype localising to the 15%-35% Percoll interface (Figure 3b). We therefore hypothesised that if *rmpA* variants were generated stochastically during non-selective growth, they could be isolated from the 15%-35% interface of WT-inoculated gradients. To investigate this, we analysed mucoidy and *rmpA* expression in ∼90 representative colonies from the top (WT HV*Kp*), and 15%-35% interface (predicted *rmpA* mutants) of gradients derived from overnight cultures of a single SGH10 WT founder colony. This analysis indicated that the top fraction of each strain was entirely resistant to sedimentation and no mutations were detected in *rmpA* (Fig 3c). Conversely, in the putative low-capsule fraction, ∼50 % of colonies were no longer mucoid, with this phenotype correlating with both frequent frameshift mutations in *rmpA* and with loss of the virulence plasmid. Similar data were obtained following analysis of gradients inoculated with strain SB4538 (data not shown). This indicates that sub-populations of *rmpA*-mutants and virulence plasmid-negative strains are generated during a single round of overnight growth in rich media. To confirm that poly-G mutations in *rmpA* drove reductions in capsule expression and mucoidy, we assayed the sedimentation resistance and uronic acid content of WT HV*Kp, rmpA*-OFF truncation variants isolated from the 15%-35% Percoll interface, and *rmpA-*OFF strains complemented with a functional copy of *rmpA* on a plasmid. This analysis confirmed that mutants were no longer mucoid (Fig 3d) and expressed less polysaccharide capsule (Fig 3e), and that both phenotypes could be rescued by complementation in trans with the WT *rmpA* gene. Interestingly, truncation of *rmpA* in the SGH10 background led to a smaller reduction of capsule expression relative to the SB4538 background (Fig S4), likely explaining why selection of *rmpA* mutants in SGH10 was weaker in the infection assays. Together, these data indicate that high frequency mutations in *rmpA* generate sub-populations of bacteria expressing different amounts of capsule in the absence of selection.

### *rmpA* mutations mediate reversible phase variation of the mucoid phenotype

Given that repetitive DNA is known to mediate reversible switches in gene expression in *Neisseria* and *Campylobacter ssp.,* we tested the hypothesis that revertant mutations in the *rmpA* poly-G repeat would mediate phase variation of the mucoid phenotype. To this end, we cultured single colonies of *rmpA*-OFF variants isolated from low density regions of a Percoll gradient overnight at 37°C in LB, before subjecting these cultures to a second round of gradient centrifugation, harvesting individual colonies from the high-density fraction, and assaying mucoidy and *rmpA* expression states as above. Imaging of gradients indicated that the density of these cultures in Percoll was readily reversible (Fig 4a). Genetic and phenotypic analysis of representative colonies indicated these populations had regained the mucoid phenotype (Figure 4b) and consisted almost exclusively of bacteria in the *rmpA*-ON state driven by reversion to the WT G10 repeat length (Figure 4c). These data indicate that ON-OFF phase variation of capsule mucoidy and hyperexpression is reversible, driven through expansion/retraction of poly-G repeats in *rmpA*.

**Figure 4.**
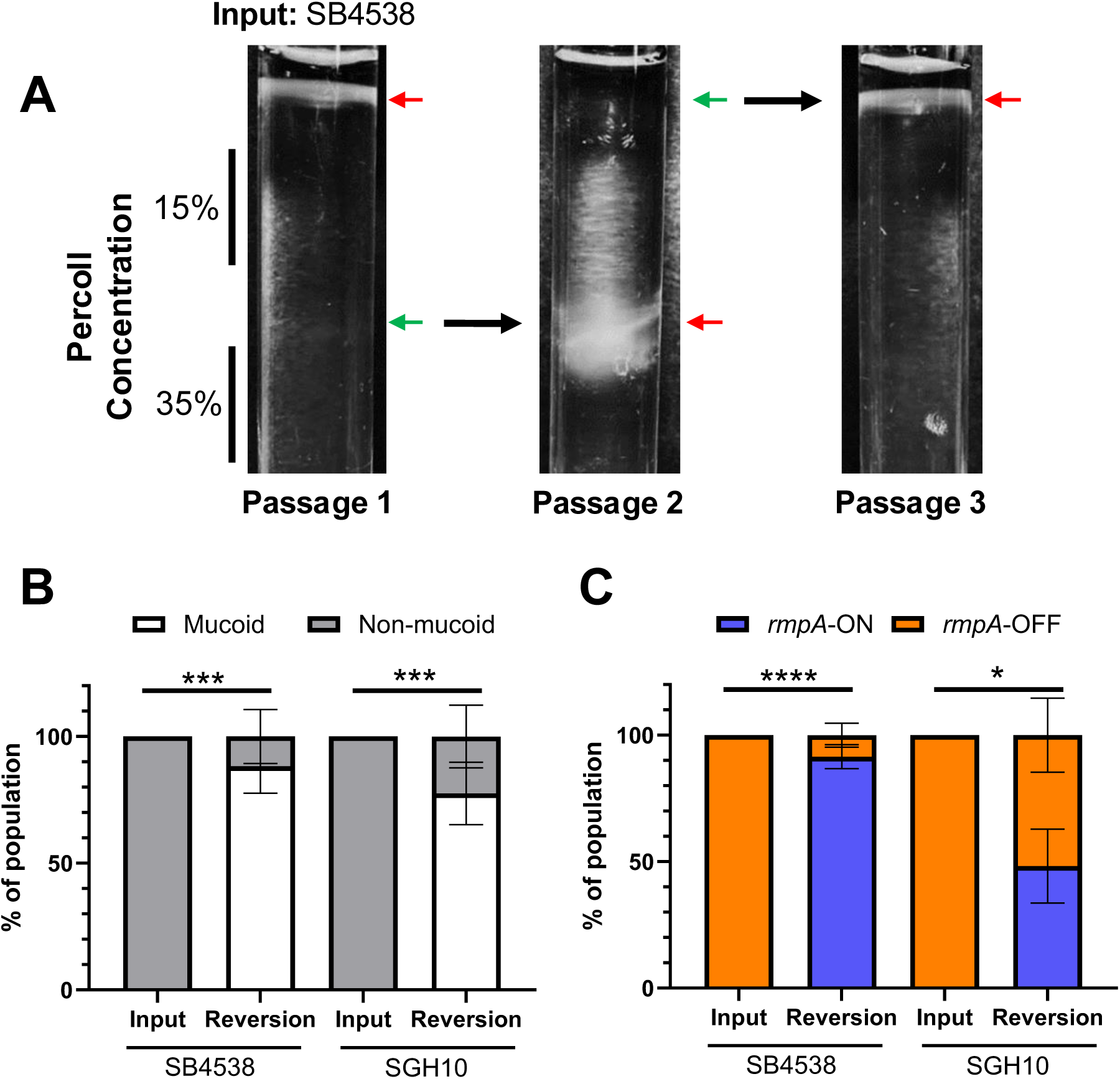
*rmpA* mutations mediate reversible phase variation of the mucoid phenotype. (A) Images of Percoll gradients following cyclical selection of cultures with high, low (15%-35% interface), and high density. Passage numbers are indicated at the bottom of the image, and subsequent passages were initiated by removing 100 µL of culture from target regions in the prior passage (indicated by green arrows), inoculating into fresh LB overnight, and re-separating on a gradient the following day. Red arrows indicate the localisation of the bacterial culture. (B) Quantification of mucoidy in multiple colonies derived from *rmpA*-OFF input populations, and revertant mucoid output populations isolated from the top fraction of Percoll gradients in both the SB4538 and SGH10 background. Error bars represent the standard deviation. White stacked bars; proportion of mucoid colonies, grey stacked bars; proportion of non-mucoid colonies. (C) Quantification of *rmpA* expression state in multiple colonies derived from *rmpA*-OFF input populations, and revertant mucoid output populations in both the SB4538 and SGH10 background. Error bars represent the standard error of the mean. Orange stacked bars; proportion of *rmpA*-ON colonies, blue stacked bars; proportion of *rmpA-*OFF colonies. Data are representative of at least 3 independent experiments, and statistical significance was determined by comparing the proportion of colonies with each phenotype/genotype using a Generalise Linear Mixed Model. ***; P>0.0005, **; P>0.005, *; P>0.05, ns; not significant.

### *rmpA* phase variation modulates virulence and serum sensitivity

Given that capsule mucoidy is an established virulence determinant in HV*Kp* ^17^, we next aimed to determine if *rmpA* phase variation would drive altered virulence in an acute murine pneumonia model. We therefore infected mice intranasally with 10^5^ CFU of WT HV*Kp* (*rmpA*-ON), an isogenic *rmpA*-OFF variant, and a *rmpA*-ON revertant strain isolated based on density in Percoll (Fig 4). We observed that all SB4538 variants had an indistinguishable capacity to colonise the lung, but that inactivation of *rmpA* by phase variation almost entirely prevented systemic dissemination to the liver and spleen (Figure 5a). In-keeping with a reduced role for *rmpA* phase variation in regulation of capsule in SGH10 (Figure S4), we observed moderate but significant decreases in the number of bacteria recovered from lungs and livers of mice infected with an SGH10 *rmpA*-OFF strain compared to *rmpA*-ON variants (Figure 5b). As capsule has been shown to reduce serum complement deposition at the bacterial surface^21^, we next tested the hypothesis that differences in susceptibility to serum complement underpinned the observed changes in the *in vivo* virulence phenotype by testing the survival of *rmpA*-ON, OFF, and revertant ON strains in naïve human serum. Consistent with the murine infection data, we observed that both the WT serotype K1 strain SGH10, and its *rmpA*-OFF, and *rmpA*-ON revertant derivatives were highly resistant to the bactericidal activity of undiluted serum (Fig 5c). Instead, whilst the WT K2 *rmpA*-ON strain exhibited resistance to the bactericidal activity of 50% serum, the *rmpA*-OFF variant was highly susceptible. Importantly, reversion of the poly-G repeat to the ON length, was able to restore resistance to serum killing (Fig 5c). Together, these data indicate that, at least in some strain backgrounds, the reversible ON/OFF phase variation of *rmpA* drives a phenotypic switch in systemic spread in mice that may, in part, be due to altered susceptibility to serum complement-mediated killing.

**Figure 5.**
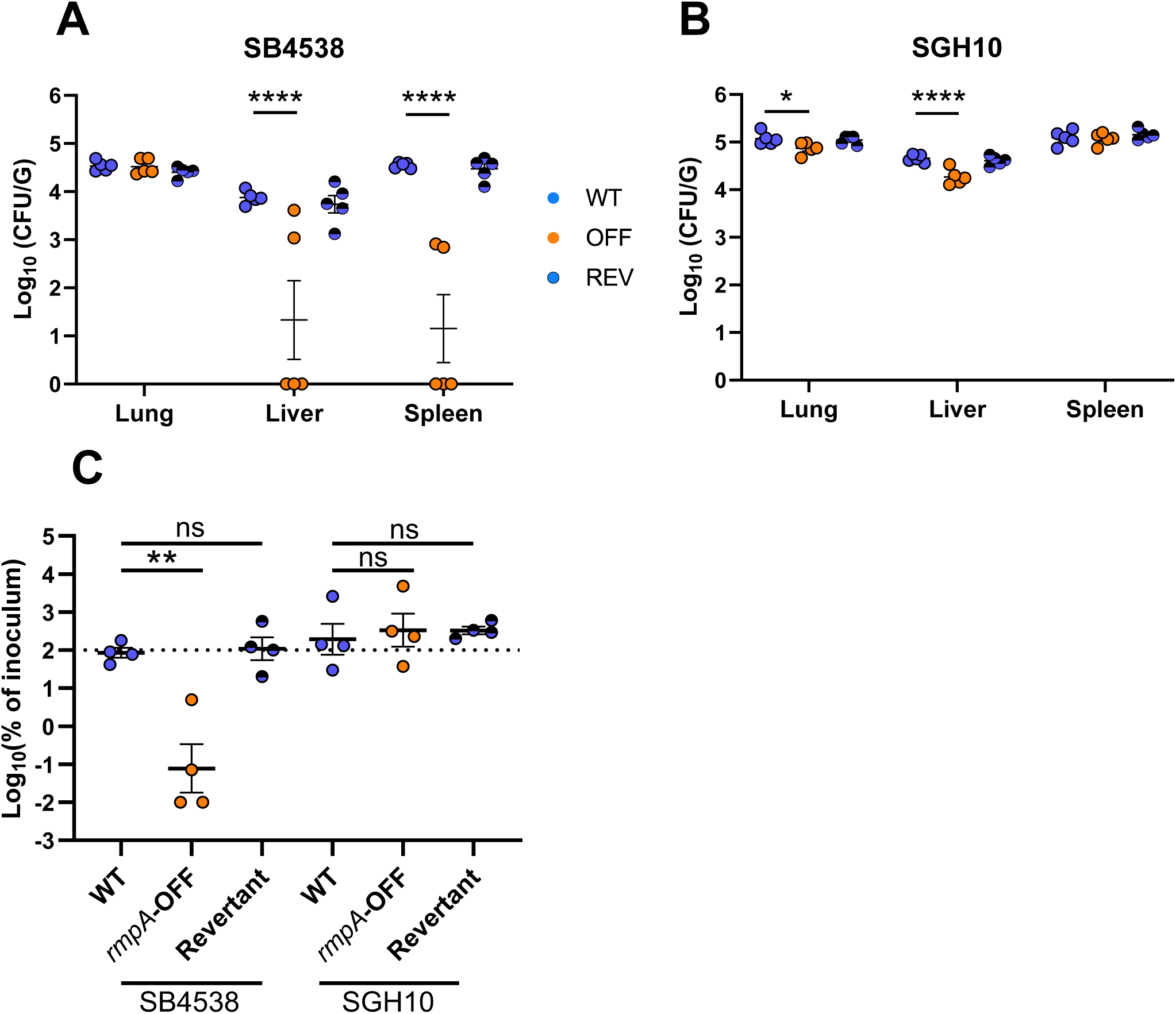
*rmpA* phase variation modulates bacterial virulence and serum sensitivity. (A-B) Bacterial counts expressed as CFU per gram of tissue, in the blood, liver, and spleen of CD1 mice following intranasal infection of *rmpA*-ON (blue), *rmpA*-OFF (orange) and revertant *rmpA* ON (blue patterned) variants in the SB4538 (A) and SGH10 (B) background. Data represents the mean and standard deviation of 5 infected animals per group. (C) Survival (blue bars) of WT and *rmpA*-OFF phase variants (orange bars) in human serum. SGH10 and its derivatives were incubated in 100% serum for 1h, whereas SB4538 derivatives were incubated in 50% serum for 1h. Control experiments demonstrating ablation of all phenotypes following heat inactivation of sera are shown in Figure S5. Data are representative of 4 independent biological replicates, and statistical significance were determined using an ordinary 1-way ANOVA. ns; non-significant, **; P>0.005.

### Repeat-mediated phase variation is a conserved feature of the *rmp* locus

The *rmp* locus is mobilised by multiple distinct mobile genetic elements (MGEs), including plasmids KpVP-1 and KpVP-2, and the integrative conjugative element ICE*Kp1* ^31^. As phase variation of mucoidy is likely to be an important adaptive strategy for HV*Kp,* we hypothesised that the repeat-mediated switching mechanism would be conserved in all *rmp* loci from these elements. To this end, we performed a comparative analysis of repeat lengths and associated ON/OFF expression states in the RmST allele database, which is derived from concatenated sequences of *rmpA, rmpD* and *rmpC* genes, and represents the full breadth of *rmp* genetic diversity in sequenced *Kp* ^31^. We first annotated a conservation map derived from alignment of the concatenated genes from each RmST lineage, with the location of homopolymeric SSRs. This map revealed, in addition to the *rmpA* poly-G (bases 276-285) repeat described so far, the presence of both a poly-T (bases 30-38) and poly-A (bases 136-144) repeat in *rmpD*, a poly-G repeat in *rmpC* (bases 325-333), and an additional poly-T (bases 63-70) repeat in *rmpA* all consisting of 9 or more monomers (Figure S6a). We chose to exclude analysis of a further polyA (bases 189-196) and polyT (bases 209-216) repeat in *rmpC,* as these tracts consisted of 8 nucleotides, which are below previously described length cutoffs for phase variable genes^51,58^. In all cases, these homopolymers mapped to areas of low sequence conservation indicating they are a major driver of genetic diversity at this locus (Figure S6b). We next quantified the lengths of each repeat and mapped this data to a phylogenetic tree annotated with their phylogenetic lineage (*rmp1-4*) and MGE (e.g. KpVP-1, KpVP-2, ICEKp1) ^31^ (Figure 6). Gene truncation alone does not imply propensity for phase variation, as multiple stop codons within the reading frame may lead to production of a constitutively non-functional protein. The hallmark of SSR-mediated PV is frameshifts in full-length gene expression due to expansion and retraction of the repeat number. To model this, we performed *in silico* expansions of repeats from each unique *rmp* locus by adding 1, 2, and 3 monomers prior to translation. The resulting protein products were expressed as a percentage of the maximum possible amino acid length to infer their predicted ON or OFF expression states (translated products >95%, or <95%, of the expected length respectively). We observed that 91.8% of *rmpA,* 91.2% of *rmpD* and 84.8% of *rmpC* gene truncations were sensitive to expansion of the poly-G, poly-A, and poly-G, repeat tracts respectively, suggesting that changes in repeat length – rather than additional mutations – were responsible for driving phase variation. Furthermore, we found that phase variation occurred frequently across the phylogeny, with extensive repeat length heterogeneity present within each individual lineage (Figure 6), indicating that phase variation was not the result of clonal expansion of fixed repeat-length alleles in specific sub-lineages, but rather an ongoing, dynamic process. Whilst most changes in expression were driven by alterations in the number of repeats, we also observed cases where the SSR was interrupted by one or more divergent nucleotides with some extreme cases resulting in complete loss of the SSR tract, indicating loss of capacity to undergo phase variation (Table 3). Importantly, these sequences were rare and arose in each lineage but showed no evidence of clonal expansion, indicating strong selection for maintenance of the switching mechanism. Analysis of RmpD and RmpC AlphaFold models for full length and truncated protein variants demonstrated loss of key functional regions, indicating that repeat expansions likely lead to loss of function (Figure S7). These data indicate that phase variable repeats are a conserved feature of all *rmp* lineages, and that both *rmpD* and *rmpC* are likely also phase variable.

**Figure 6.**
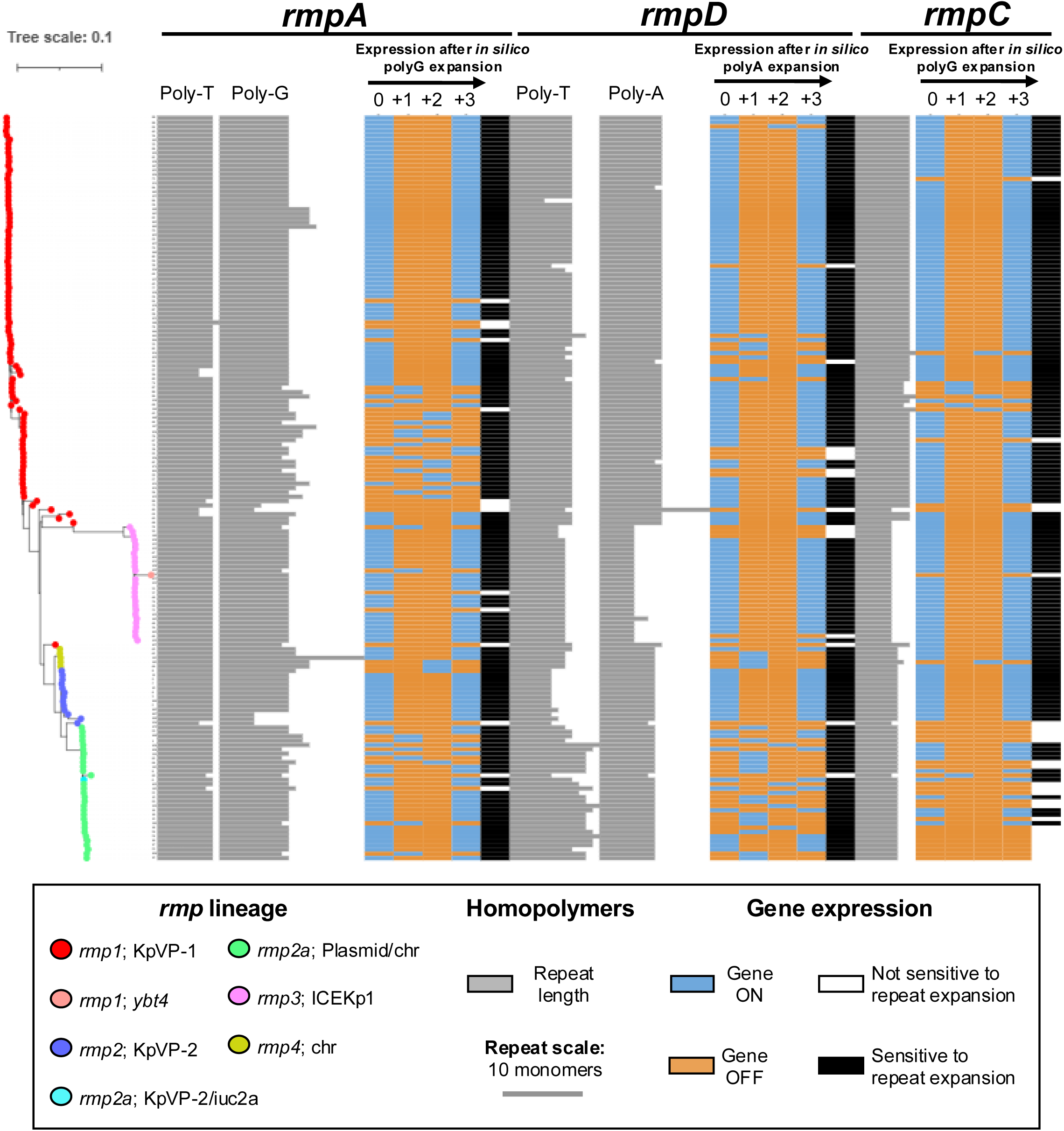
Repeat-mediated phase variation is a conserved feature of the *rmp* locus. (A) Maximum likelihood phylogenetic tree of *rmp* loci outlined in the RmST typing scheme with 100 bootstrap replicates. Nodes are colored according to their associated mobile element and are indicated in the legend. Grey bars indicate the length of the homopolymeric repeat in each gene. Heatmaps indicate the ON/OFF expression of each gene in the locus with its native repeat length (0) and following *in silico* expansion of the repeat by 1, 2, or 3 nucleotides. The capacity of sequences to alternate between ON and OFF expression states following *in silico* expansion of the indicated repeat are shown by black bars. ON/OFF expression states are derived through translation of the DNA sequences, and sequences producing a protein length >95% of the maximum were coded ON, while all other sequences were coded OFF.

**Table 3.**
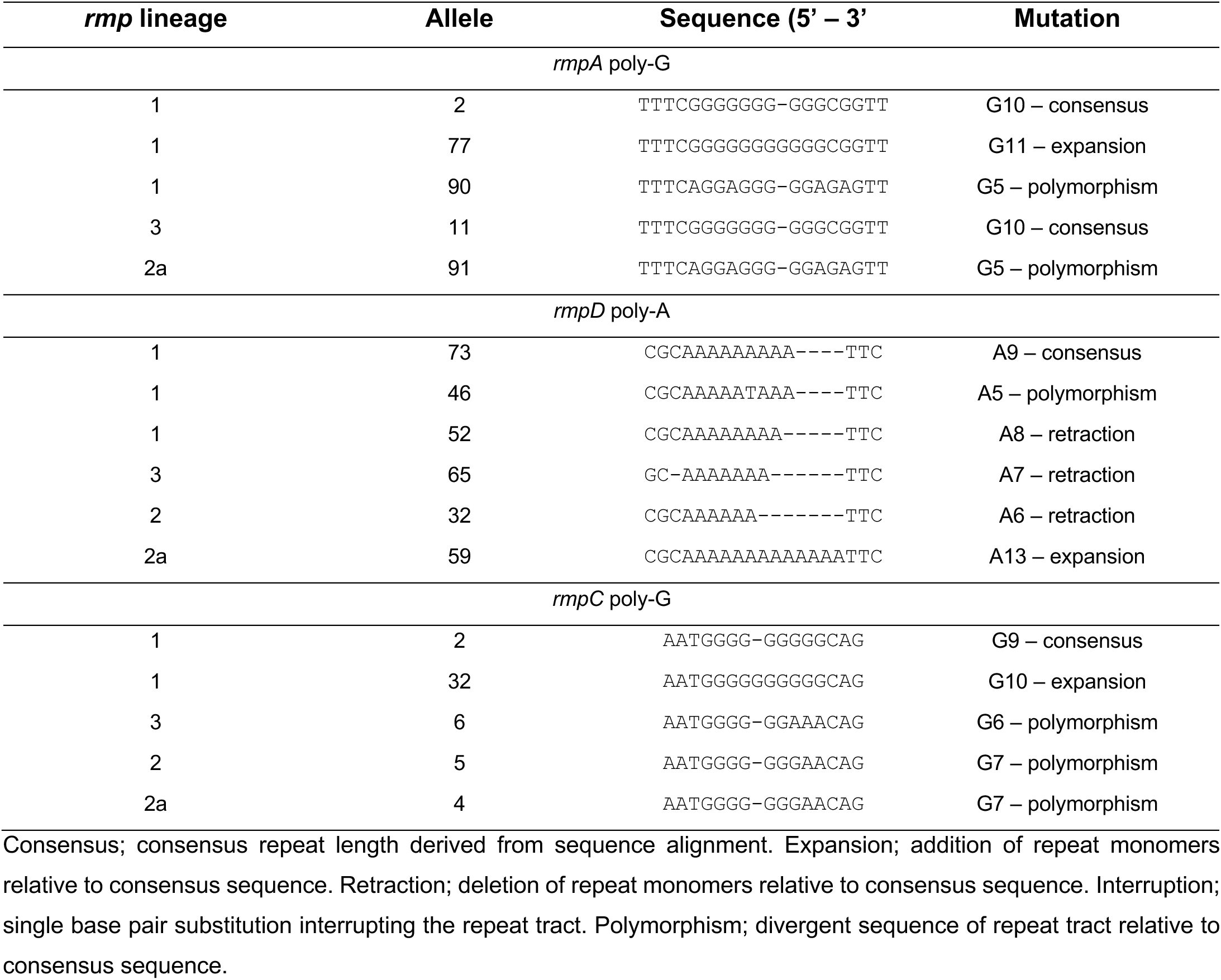
Representative polymorphisms at homopolymeric repeats in the *rmp* locus.

### Independent repeat mutations drive lineage-specific, combinatorial expression of capsule and mucoidy

Mutation rates of DNA repeats increase as a function of repeat length ^59^. This led us to hypothesize that selection for different switching frequencies may drive lineage-specific variations in repeat number or type. We therefore compared *rmp* repeat lengths in the RmST phylogenetic lineages with greater than 10 representative sequences (*rmp1;* KpVP-1, *rmp2;* KpVP-2, *rmp2a;* plasmid/chromosome, and *rmp3;* ICE*Kp*1; Figure 7a). The *rmpA* poly-G repeat showed the same modal length (G10) in all lineages, suggesting selection for this as an optimal length. In contrast, repeat tracts in *rmpD* and *rmpC* had distinct modal lengths in each lineage (Figure 7a), suggesting ongoing selection for mutation rates associated with different repeat lengths. Comparison of Alphafold models for RmpA containing either a G10 or G20 repeat, respectively, showed no obvious difference in predicted protein structure, indicating that changes in repeat length likely do not confer changes in protein function (Figure 7b). These data indicate that all lineages of *rmp* show heterogenous repeat lengths evident of different switching rates, and that repeat length likely has little impact on protein function.

**Figure 7.**
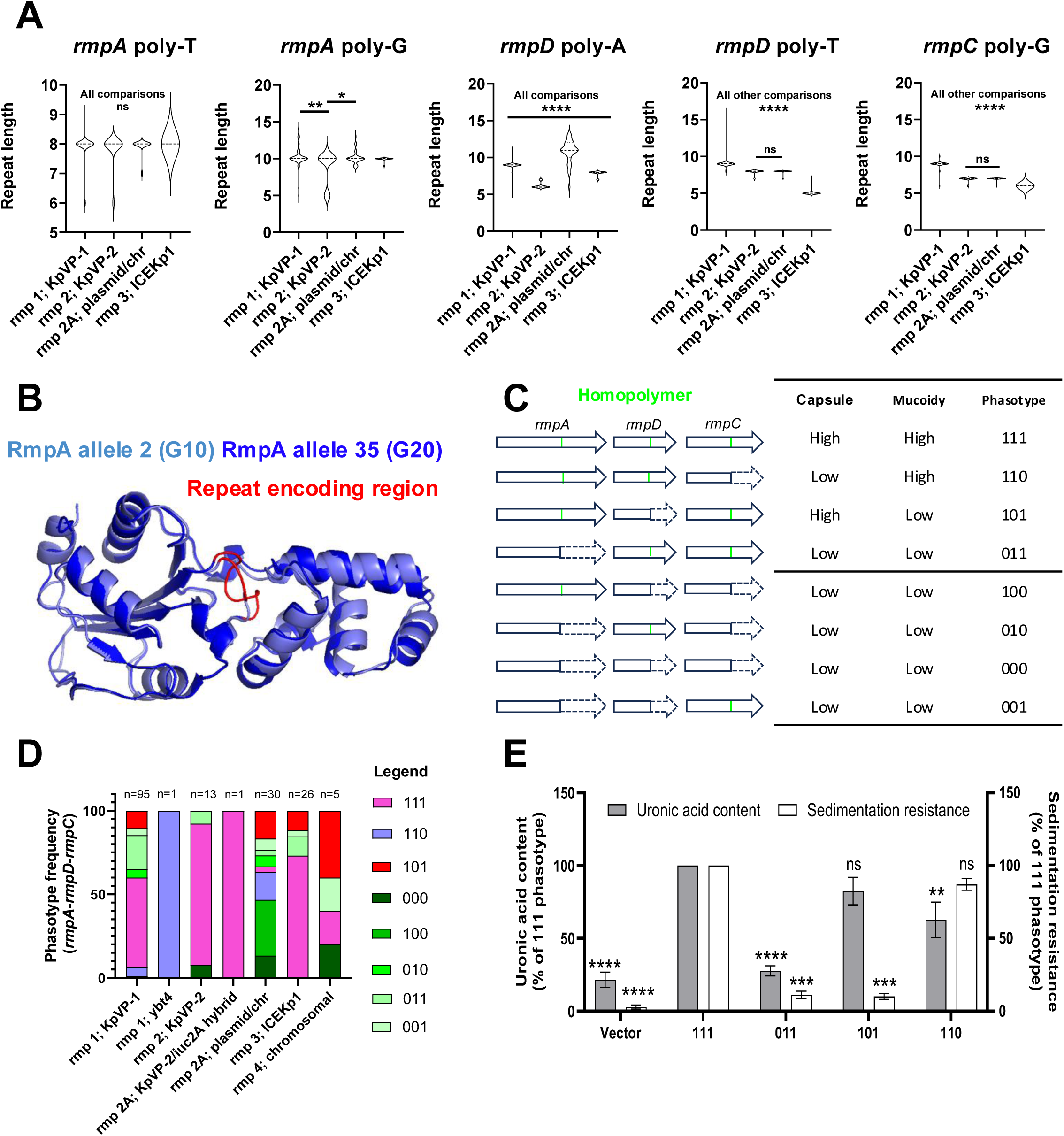
Independent repeat mutations drive lineage-specific, combinatorial expression of capsule and mucoidy. (A) Violin plot depicting length of homopolymeric repeats in each gene of the *rmp* locus from lineages *rmp1* (KpVP-1; n=95), *rmp2* (KpVP-2; n=13), *rmp2a* (plasmid/chr; n=30), and *rmp3* (ICEKp1; n=26). Statistical significance was determined by one way ANOVA. ***; P>0.0005, **; P>0.005, *; P>0.05, ns; not significant. (B) Alignment of alphafold structures of RmpA derived from *rmpA* allele 2 (dark blue; G10 length) and allele 35 (light blue; G20). The repeat encoding region is shown in red. (C) Schematic depicting truncated *rmp* loci and their predicted capsule and mucoid phenotypes. (D) Distribution of combinatorial gene expression phasotypes in each major *rmp* lineage. Phasotypes are coloured according to the legend, and the number of sequences represented are shown above each bar. (E) Analysis of capsule mucoidy (sedimentation resistance; white bars) and uronic acid content (grey bars) in a SB4538ΔpVir strain transformed with the pBR322 empty vector, and constructs harbouring phasotype variants 111, 011, 101, 110 or *rmp* with the native promoter. All data are expressed as a percentage of the 111 phasotype strain. Statistical significance were determined by one-way ANOVA with comparison to the 111 phasotype control. Dat are representative of 6 independent experiments. ns; non-significant, **; P>0.005, ***; P>0.0005, ****; P>0.00005.

Given the distinct roles of *rmpA* as an autoregulator of the operon, *rmpD* as a regulator of capsule polymer length and mucoidy, and *rmpC* as a positive regulator of the capsule biosynthesis operon, we hypothesised that independent mutations in each gene would give rise to 4 different combinatorial capsule phenotypes, ranging from low mucoidy and low capsule expression, to high mucoidy and high capsule expression. We therefore assigned combinatorial *rmpA-rmpD-rmpC* expression states, termed ‘phasotypes’, to each *rmp* locus variant, scoring the binary ON/OFF expression of each gene in the locus as 1 and 0, respectively (Figure 7c). For example, a locus with all genes switched ON is scored 111, whereas a locus with all switched OFF is scored 000. Extending this analysis to the RmST database indicated a spread of predicted phasotypes across all lineages. 6/8 and 4/8 phasotypes were detected in *rmp1* and *rmp3,* respectively, with the 111 phasotype being the most common in both (Figure 7d). In contrast, in *rmp2a,* the 111 phasotype was observed in a minority of sequences, instead being dominated by phasotypes harbouring frequent truncation in *rmpA, rmpD* and *rmpC* (Figure 7d). These data suggest that independent truncations in *rmpA, rmpD,* and *rmpC* generate combinatorial patterns of gene expression, and that these profiles vary by *rmp* lineage.

To investigate the relationship between *rmp* phasotype and capsule phenotype, we expressed different WT and repeat-truncated *rmp1* variants under the control of their native promoter from the medium-copy number vector pBR322 in a variant of SB4538 cured of the large virulence plasmid, and assayed mucoidy and uronic acid concentrations in the recombinants (Figure 7e). To avoid confounding effects of sequence variation elsewhere in the locus, we assayed isogenic copies of *rmp1* (identical sequence to strain SB4538) with changes solely in the *rmpA* poly-G (G10 to G9; 011 phasotype), the *rmpD* poly-T (T9 to T8; 101 phasotype), and the *rmpC* poly-G (G9 to G8; 110 phasotype). This analysis revealed that expression in trans of a functional *rmp1* locus significantly increased expression and mucoidy of the capsule relative to the empty vector control. Strikingly, repeat retraction-mediated truncation of *rmpA* abolished both phenotypes, while truncation of *rmpD,* and *rmpC*, abolished only capsule mucoidy or hyperexpression, respectively, indicating that repeats in each gene mediate independent ON/OFF switches in these two properties of the capsule. Together, these data demonstrate independent mutations at DNA repeats in each gene of the *rmp* locus, generate combinatorial patterns of capsule hyperexpression and mucoidy.

## Discussion

HV*Kp* express a thick and physically mucoid capsule, driven by the MGE-encoded *rmp* locus, that promotes their virulence ^18,19,60–63^. Our results demonstrate that the mucoid phenotype is prone to reversible ON/OFF phase variation allowing HV*Kp* to alternate between phenotypic states favouring either host cell infection or resistance to serum bactericidal killing. Since its first description in 1986 in Taiwan, the HV*Kp* pathotype has disseminated across the globe ^3^. Following initial episodes of asymptomatic gut colonisation, HV*Kp* transition to the bloodstream, and disseminate to systemic tissues where bacterial cells occupy both intra and extracellular spaces ^1,4,26,28,64–68^. This diversity of niches would appear at odds with constitutive high level capsule expression, given that mucoid capsule is required for survival in serum ^21,69^ but significantly hinders interaction with and invasion of host cells ^29,56,70^. Our work provides the first evidence of a mechanistic basis for this paradox, revealing that *rmpA* phase variation directly regulates mucoid capsule expression (Fig 3) and allows HV*Kp* to alternate between states of host cell colonisation (Fig 1) and resistance to serum bactericidal killing (Fig 5). Our study demonstrates that phase variation drives population heterogeneity in capsule expression in the absence of selection, indicating that *rmpA* is a “contingency locus” – a region of the genome prone to localised hypermutation to generate ‘pre-emptive’ genetic and phenotypic heterogeneity to allow rapid adaptation to fluctuating selective pressures. As capsule and mucoidy are critical determinants of HV*Kp* virulence, our findings markedly alter our view of how this pathogen disseminates both within and between hosts.

HV*Kp* are highly diverse, consisting of multiple serotypes, phylogenetic lineages, and differing repertoires of virulence genes. Recent studies have demonstrated that the *rmp* locus was the major contributor to hypervirulence in murine infection models across multiple HV*Kp* clonal lineages,^71,72^ and that carriage of *rmpA* and *rmpA2*, in addition to *iroB*, *iucA*, and *peg-344*, were significantly associated with poor patient outcome in humans.^73^ These studies highlight the importance of RmpA as a driver of human disease. Notably, the *rmp* locus consists of at least 7 distinct lineages that are all able to drive capsule hyperexpression and mucoidy in HV*Kp* ^31^. Though each of these distinct lineages is mobilised by different virulence plasmids and integrative conjugative elements, our results reveal that all *rmp* loci contain several homopolymeric tracts suggestive that phase variation is a conserved feature of all HV*Kp.* Interestingly, repeat lengths varied across *rmp* loci harboured by different MGEs, with longer repeats in loci associated with the widely disseminated KpVP-1. As studies in *Campylobacter*, *Neisseria*, and *Haemophilus* have indicated that mutation rates increase as a function of repeat number ^74–77^, it is tempting to speculate the success of KpVP-1 is due in part to higher rates of associated *rmp* phase variation. Our work therefore mandates further characterisation of the strength of interaction between *rmp* and different HV*Kp* capsule type and how phase variation rates differ across HV*Kp*, in both experimental virulence models and during genomic surveillance. We have further demonstrated that independent DNA repeats in *rmpA, rmpD,* and *rmpC* drive 8 combinatorial ON/OFF genotypes termed ‘phasotypes’, four of which we have functionally validated. Given that recent studies have shown that capsule mucoidy and hyperexpression drive distinct immune evasion programs and interactions with the host ^37,38^, we suggest that the combinatorial nature of *rmp* phase variation, provides a ‘swiss-army knife’ of capsule phenotypes to allow HV*Kp* to adapt to distinct selection pressures in the host.

Expression of the HV*Kp* capsule is known to be controlled through the activity of transcriptional regulators which respond to the host environment signals ^30,39,40,42,78–80^. Whilst undoubtably required for pathogen fitness during prolonged colonisation, such transcriptional remodelling may be insufficient for the pathogen to adapt to rapid changes in environment such as those encountered during dissemination within and between hosts. Our data support a model in which the rapid generation of variants in the absence of selection through *rmpA* phase variation pre-adapt populations of HV*Kp* to changes in environment associated with different requirements for capsule expression. Further supporting our hypothesis that mutations in capsule regulators are a key feature of *Kp* host adaptation, Unverdorben *et al* (2025) recently demonstrated that gut colonisation in mice strongly selects for capsule-inactivating mutations in an ST258 strain.^81^ Ernst *et al.* (2020) found that similar capsule-inactivating mutations enable ST258 strains to adopt persistent or systemic infection programs.^29^ Supporting a key role of repetitive DNA in host adaptation, Teng *et al.* recently demonstrated that expansion of a poly-T repeat in the *rmp* promoter region modulates mucoidy in an ST11 strain ^82^. Song *et al.* ^83^ also demonstrated that both an insertion element ISKpn26 upstream of the *rmp* locus and mutations in the *rmpA* poly-G repeat mediated reduction in mucoidy, driving persistent infection in a patient with a scrotal abscess complicated by UTI, indicating that mutations at this locus have clear clinical relevance. The rapid, reversible nature of the mechanism we have uncovered indicates that rather than simply representing a fixed loss of function, mutations at the *rmpA* poly-G repeat instead enable rapid alternation distinct infection programs. Taken together our work and these prior studies, support the sampling of multiple bacterial isolates from single patients to shed light on how genetic heterogeneity alters disease pathogenesis and may inform clinical management.

The emergence of convergent MDR HV*Kp* is an urgent threat. Capsule expression is a major barrier to horizontal gene transfer and acquisition of foreign DNA is rare in HV*Kp* compared to classical *Kp* ^84,85^. Despite this, there have been reports of acquisition of MDR plasmids by HV sequence types (STs), and conjugative transfer of the large virulence plasmid from HV STs to classical STs ^11,86–88^. As a hypermucoid capsule limits horizontal gene transfer, we postulate that phase variable capsule expression may enable uptake and/or transfer of such mobile elements with only temporary reductions in capsule expression and mucoidy, mitigating the immediate fitness cost of the convergent phenotype. In the future, further functional analysis of capsule phase variation across all HV*Kp* will shed light on the evolutionary dynamics shaping the spread of hypervirulence and drug resistance.

Lastly, capsular polysaccharide is a lead antigen for vaccines in pre-clinical development against HV*Kp* ^89,90^. Capsule phase variation will not undermine the efficacy of such vaccines as the capsule is essential for virulence but may hold implications for the exposure of sub-capsular protein antigens for use in novel vaccine formulations to afford more broad strain coverage^91^. Our work therefore warrants further investigation of the interaction of capsule phase variation with access of antibodies to the bacterial surface.

In conclusion, we have established a mechanism driving reversible phase variation of the mucoid phenotype in HV*Kp*. Expansion and retraction of repetitive DNA in *rmpA* alters capsule phenotypes and provides opportunities to rapidly adapt to extracellular and intracellular environments during infection. Our work highlights that therapeutic strategies against HV*Kp* infections likely need to dual target the capsule and antigenic structures underneath the capsule.

## Acknowledgements

We would like to acknowledge members of the Odendall, Hill, and Torraca labs for their feedback through this project. We would also like to thank Professor Jeremy Brown, and Professor Chris Bayliss for useful advice during this project and critical reading of the manuscript. We thank Dr Julien Bergeron for assistance with generating Alphafold structures. We thank Dominique Decré and Sylvain Brisse for kindly providing strain SB4538. J.J.W is funded by Professor Anthony Mellow’s Charitable Settlement and the Wellcome Trust (grant 320977/Z/24/Z). A.J.M is funded by the Wellcome Trust (grant 225315/Z/22/Z).

## Materials and Methods

### Bacterial strains, genetic manipulation, and culture conditions

The serotype K1, Sequence Type (ST) 23 strain SGH10 was obtained from ATCC, whilst the K2, ST25 strain SB4538 was kindly gifted by Professor Sylvain Brisse. Strains were routinely cultured on LB (Miller) agar plates (Merck Millipore; 110283) supplemented with antibiotic selection where required. Strains were cultured in liquid LB (Merck Millipore; 110285) at 37°C with shaking (200 rpm) overnight and were routinely sub-cultured 1:100 the following day and grown until mid-logarithmic phase before centrifugation and resuspension in PBS (Gibco; 18912.014) for inoculation in assays. Electrocompetent *Kp* strains were generated as previously described ^92^, and transformations were performed using a Biorad Micropulser. All constructs used in this study were either synthesised and sub-cloned by BioBasic or were kindly provided by collaborators. All bacterial strains and plasmids are described in Table 1.

### Whole genome sequencing and variant calling

To detect mutations in cell-associated *Kp,* we subjected individual clones to short read illumina whole genome sequencing by MicrobesNG (Birmingham, UK). Briefly, single bacterial colonies were inoculated into fresh LB and grown until the mid-log growth phase before being resuspended in DNA/RNA Shield (Zymo Research, USA). Genomic DNA libraries were prepared using the Nextera XT Library Prep Kit (Illumina, San Diego, USA) following the manufacturer’s protocol but with a 2-fold increase in input DNA, and a PCR elongation time of 45 seconds. DNA quantification and library preparation were carried out on a Hamilton Microlab STAR automated liquid handling system (Hamilton Bonaduz AG, Switzerland). Libraries were sequenced on an lllumina NovaSeq 6000 (Illumina, San Diego, USA) using a 250 bp paired end protocol. Raw reads were adapter trimmed using Trimmomatic version 0.30 with a sliding window quality cutoff of Q15^93^. De novo assembly was performed using SPAdes version 3.7^94^, and contigs were annotated using Prokka 1.11^95^. Candidate virulence plasmid-negative mutants were subject to further long read whole genome sequencing by MicrobesNG (UK). DNA libraries were generated using the Oxford Nanopore Technologies (ONT) SQK-RBK114.96 kit (ONT, United Kingdom) from 200-400 ng of High Molecular weight (HMW) DNA and was sequenced on a or FLO-MIN111 (R10.3) flow cell in a GridION (ONT, United Kingdom). Genome assembly was performed using Unicycler version 0.4.0, and contigs are annotated using Prokka version 1.11. Loss of the virulence plasmid and virulence gene content in genome assemblies was determined using Kleborate^96^. For variant calling, reads were aligned to their respective WT sequence using BWA mem v0.7.17 and were subsequently processed using SAMtools v1.9. Variants were called using VarScan v2.4.0 with considering variant allele frequencies greater than 90%. Effects of the variants were predicted and annotated using SnpEff v4.3.

### Mammalian cell culture and *in vitro* infections

THP-1 cells were maintained in RPMI supplemented with 10% FCS and Glutamax (Gibco; 61870036). For differentiation, cells were seeded in 24 well TC-treated plates (Corning; 10380932) at a density of 3.5 x 10^5^ cells per well in the presence of 100 ng/mL PMA (Merck Millipore; P1585). After 2 days, the media was changed to complete medium without PMA and cells were rested for an additional day prior to infection. For infection, overnight cultures were sub-cultured to mid-logarithmic phase before infection of cells at an MOI of 100. Plates were synchronised by centrifugation at 200xG for 5 minutes, and cells were left infected for 30 minutes. At this stage, some plates were washed 3x in PBS, lysed in 0.5% saponin, serially diluted, and plated to measure bacterial adhesion, whereas the others were treated with 100 ug/mL gentamicin in cell culture medium for 30 minutes to kill extracellular bacteria. Wells were subsequently washed, lysed, and plated as above at the required time point. A549 cells were routinely cultured in DMEM + glutamax (Gibco; 10566), were passaged at ∼80% confluence, and were seeded for infections at a density of 1 x 10^5^ cells per well in 24-well plates. Infections were performed as above, but with 1h of initial infection followed by 1h of gentamicin treatment.

### Bacterial survival in normal human serum

Pooled normal human serum (NHS) samples were generated from 4 independent mixed sex donors through the KCL Biobank (REC reference: 19/SC/0232). Briefly, whole blood was collected by a trained phlebotomist in yellow cap BD Vacutainer collection tubes (12927696) and allowed to clot for 1h at room temperature. After centrifugation according to the manufacturer’s instructions, serum samples were taken from the top phase, pooled at a 1:1 ratio between donors, and were immediately frozen at –80°C as individual 1mL aliquots. Samples were thawed on ice immediately prior to experiments. Mid-log bacterial cultures were inoculated into 100% NHS (strain SGH10), or 50% NHS diluted in PBS (strain SB4538), to an initial concentration of ∼10^5^ CFU/mL. Heat inactivation of serum at 56°C for 1h prior to bacterial inoculation was used as a negative control for complement activity. The serum resistant serotype K2 strain Kp43816 and its isogenic, serum sensitive capsule and O-antigen mutant double mutant (Δ*manC*Δ*glf*) were used as positive and negative controls, respectively (Fig S5). Bacterial survival was monitored at 0, 1, and 3h post inoculation by quantitative plating, and data were normalised to the 0h HIA sample of each strain.

### Murine infection models

The virulence of WT HV*Kp* and their isogenic *rmpA* phase variants were determined using a murine intranasal infection model. The study was carried out using 8–10-week-old female CD1 outbred mice, obtained from Charles River, UK. Bacterial strains were grown in liquid LB at 37°C overnight and were sub-cultured 1:100 the following day and grown until late-logarithmic phase before centrifugation and resuspension in PBS. The cultures were then diluted with PBS to the desired concentration prior to infection. Before bacterial inoculation, mice were lightly anaesthetized with 2.5% (v/v) isoflurane over oxygen (1.4–1.6 litres/min). Mice were intranasally administered with approximately 10^5^ CFU/mouse in 50 µl sterile PBS. At 24 hours post-infection, mice were sacrificed, and the bacterial load in tissue homogenates (lung, liver, and spleen) were measured by colony counting.

### Ethics statement

Mouse studies were performed at the University of Leicester under project (permit no. PP0757060) and personal (permit no. I4FF857C1) licenses according to the United Kingdom Home Office guidelines under the Animals Scientific Procedures Act 1986, and the University of Leicester ethics committee approval. Where mentioned, the procedure was carried out under anaesthetic with isoflurane. Animals were observed in individually ventilated cages in a controlled environment and were monitored after infection to reduce suffering.

### Phylogenetic analysis of *rmp* loci and homopolymeric repeat length quantification

Alignments of unique *rmpA, rmpD,* and *rmpC* alleles were performed using ClustalOmega on concatenated *rmpADC* sequences as outlined in the RmST typing scheme (sequences extracted 4/06/2024) ^31^. Hotspots of variation in the alignment were visualised relative to the consensus sequence using Genious Prime. Repeat lengths in FASTA files containing concatenated *rmpA, rmpD,* and *rmpC* alleles were determined using a custom script in Rstudio (V.4.4.0) which counted the maximum number of successive C, A, T, and G nucleotides (see supplementary files). Briefly, FASTA files containing individual sequences for unique *rmpA, rmpD*, and *rmpC* alleles were read using the readDNAStringSet() function from the Biostrings package. Each DNA sequence was converted to a character string, and regular expression matching (gregexpr) was used to detect runs of adjacent, uninterrupted repeat motifs. Manual inspection of sequence alignments was performed to confirm these values corresponded to the intended repeat-containing region. Alignments of concatenated *rmp* loci were used to construct a maximum likelihood phylogenetic tree with 100 bootstrap replicates using RAXML ^97^. Phylogenetic trees were visualised and annotated with the interaction Tree of Life (ITOL) ^98^.

### Assignment of putative phase variable expression states

Assignment of putative expression states was performed using R following 3 rounds of *in silico* repeat tract expansion using to determine propensity of repeats to mediate ON/OFF phase variation. Briefly, the regions containing repeat tracts were identified as above, expanded by a single nucleotide, complete sequences were translated, and protein lengths were expressed as a % of the total length using Rstudio (V.4.4.0; supplementary files). Proteins were coded ON (1; >95% of total length) or OFF (0; <95% of total length) for downstream analysis. Combinatorial *rmp* expression states (phasotypes) were assigned based on the expression of the predicted ON (1) / OFF (0) expression states of *rmpA*, *rmpD*, and *rmpC*. For example, if *rmpA* was predicted ON, and the other two genes OFF, the phasotype was denoted ‘100’.

### Separation of bacterial cultures by Percoll gradient centrifugation

Discontinuous Percoll gradients were used to separate bacterial populations based on density as previously described ^41^. Briefly, 2mL of 50, 30, and 15% Percoll solutions (Sigma Aldrich; P1644) diluted in PBS were careful layered in 15mL Falcon tubes using a sterile Pasteur pipette. 200uL of overnight bacterial culture was gently added to the top of the gradient, before centrifugation at 3,000 xG for 30 minutes with no braking. Layers were subsequently harvested, serially diluted and plated on LB agar plates for downstream analysis. Selection of *rmpA*-ON phenotypic revertants was achieved by culturing *rmpA*-OFF mutants as above, before separating them across a 6mL 15% Percoll solution. Following centrifugation, 200uL of the upper fraction was harvested and used to inoculate 5mL of fresh LB medium. Cultures were incubated overnight, re-separated on gradients, and harvested when a visible band was present at the upper fraction. Control tubes derived from inoculation of uninfected Percoll gradients were used as controls for contamination.

### Sedimentation resistance assays

Sedimentation resistance assays were conducted by inoculating 5mL of LB broth and incubating at 37 °C overnight. The following morning, the optical density of the culture was recorded, and 1mL of culture was subject to low-speed centrifugation at 1000xg for 5 minutes. 100 uL of supernatant was subsequently taken, diluted 1:10 and the optical density recorded. Data are expressed as the ratio of the supernatant OD to the pre-centrifugation OD. High throughput centrifugation assays were conducted by inoculating a single colony into 200uL of LB broth in round bottom 96 well plates (Helena Laboratories; 92697T) and incubating statically at 37°C overnight. The following day, cultures were homogenised by pipetting, before reading the OD_600_ in a microplate reader (Perkin Elmer). The plate was subsequently centrifuged as above, before transferring 50uL of supernatant to a fresh plate, reading the OD, and analysing as above. Pellet formation in the bottom of the well was also imaged and correlated perfectly with OD_600nm_ ratios. In all experiments, mucoid and non-mucoid control strains were included for normalisation.

### Multiplex genescan analysis of phase variable gene expression states in colony lysates

To determine repeat tract lengths and putative expression states of genes of interest, we adapted a previously described method based on multiplex amplification of repeat-containing regions, fragment size analysis, and downstream bioinformatic analysis (^76^; Figure S2). Colonies were first inoculated into 200uL of LB in 96 well plates and grown statically overnight at 37°C. The following day, samples were homogenised by pipetting, and low speed centrifugation assays were performed as above. Samples were then returned to the culture plate, homogenised by pipetting, and 10uL of culture was diluted into 90uL of dH20. Sterile glycerol was added to the remaining culture to a final concentration of 10% before freezing at –80 to store variants for phenotypic analysis. The diluted culture was boiled for 10 mins at 100°C, pelleted by centrifugation at 16000xg for 10 minutes, before the supernatant containing bacterial DNA was transferred to a fresh plate. Multiplex PCR reactions targeting *rmpA, rmpC,* and *wzi* were conducted using primers (Integrated DNA Technologies) listed in Table 4, with a FAM-labelled forward primer in each case. PCR was performed with OneTaq polymerase (New England Biolabs; M0480S) according to the manufacturer’s instructions with a 58°C with a 60 second extension time. Samples were subsequently subject to an A-tailing reaction as previously described ^76^, before fragment size analysis on an ABI3730xl analyser using at LIZ1200 ladder by Azenta/Genewiz. Fragment size analysis data was exported using the thermofisher cloud platform, and peak sizes were processed with PSanalyse as previously described to extract repeat tract lengths and putative expression states ^76^. Colonies were subsequently coded *rmpA*-ON (WT fragment length length), *rmpA*-OFF (altered amplicon length due to repeat expansion/contraction) or *rmpA*-LOST (wzi positive and *rmpAC* negative). DNA in the same plate of known fragment length confirmed by Sanger sequencing was included in each run.

**Table 4.**
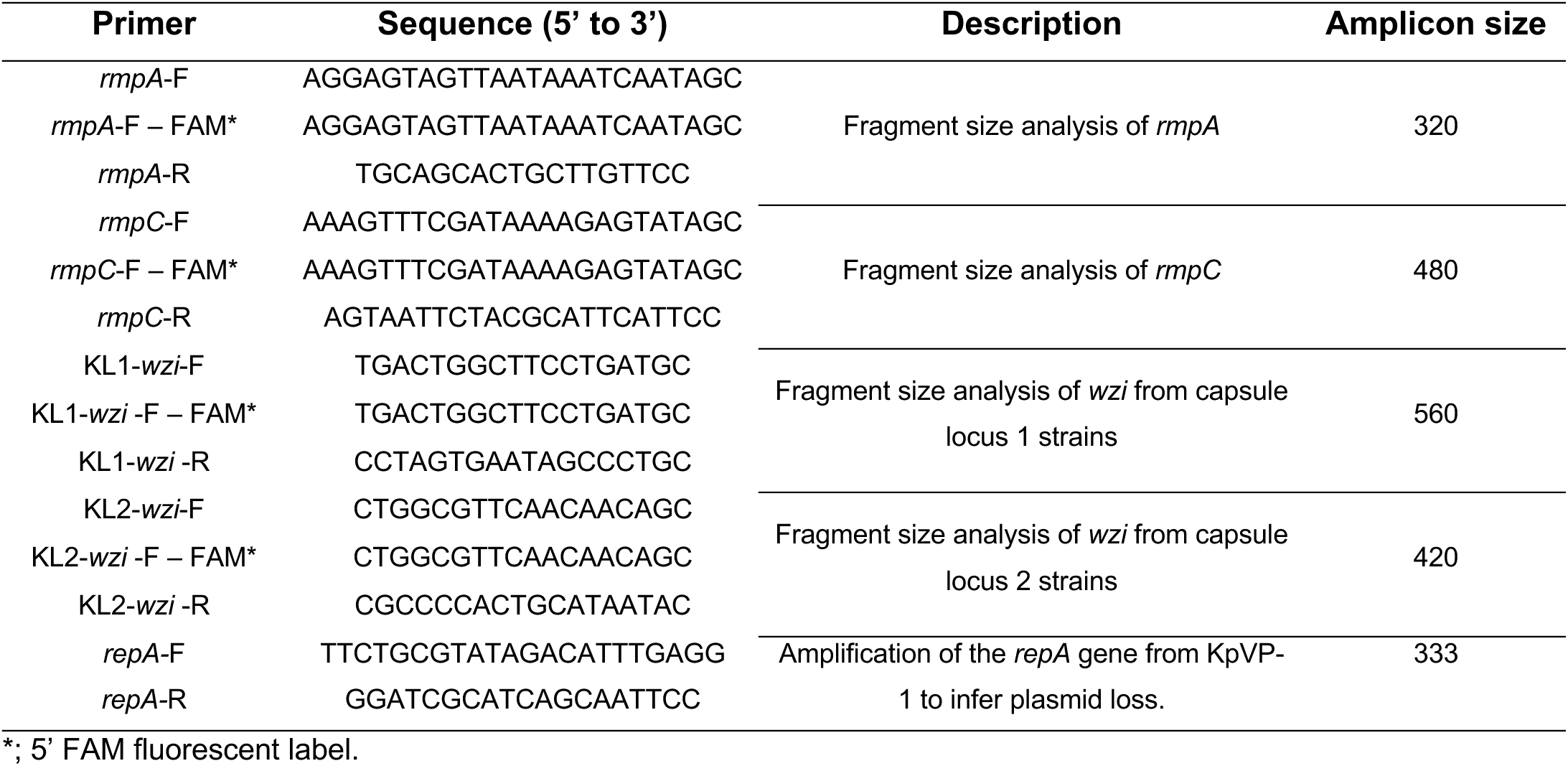
Primers used in this study.

### Uronic acid quantification

The uronic acid content of bacterial cultures were determine as previously described ^99^. Briefly, capsular polysaccharide was extracted by heating 250uL of overnight cultures with 50uL of Zwittergent 3-10 (Merck Millipore; 693021) at 50°C for 5 minutes. After centrifugation at 17,000xG, capsule was precipitated by addition of 100 uL to 400 uL ice cold absolute ethanol for 20 minutes. Capsule was subsequently pelleted by centrifugation at 17000xG for 10 minutes, dried at room temperature and resuspended in water. 1.2mL of 0.0125M sodium tetraborate (VWR International) in concentrated sulfuric acid (Supelco; 5.43827.0150) was added to the capsule preparations before boiling at 100°C. After cooling to room temperature, 10 uL of 0.3%% phenol-phenol (ThermoFisher Scientific; 417660050) was added, and absorbance was measured at 520nm. Concentrations were determined by comparison to a glucuronic acid standard curve (Merck Millipore; 843961).

### Statistical analysis

To assess changes in the proportion of mucoid versus non-mucoid colonies, and *rmpA*-ON vs *rmpA*-OFF genotypes across experimental conditions, we employed a generalized linear mixed model (GLMM) with a binomial error distribution. The model was implemented in Rstudio (v4.4.0) using the lme4 package. The response variable was the count of each genotype/phenotype, modelled as a two-column matrix using the ‘cbind’ function. Post-hoc pairwise comparisons between conditions were conducted using the emmeans package. All other statistical analysis was performed in GraphPad Prism (Version 10). Comparisons between 2 groups were performed using a student’s T-test, whereas comparisons between multiple groups were performed with an ordinary 1-way ANOVA with Dunnett’s post-hoc test for multiple comparisons. N numbers, and P values are reported in the legend for the requisite figure.

### Supplementary figure legends

**Figure S1.**
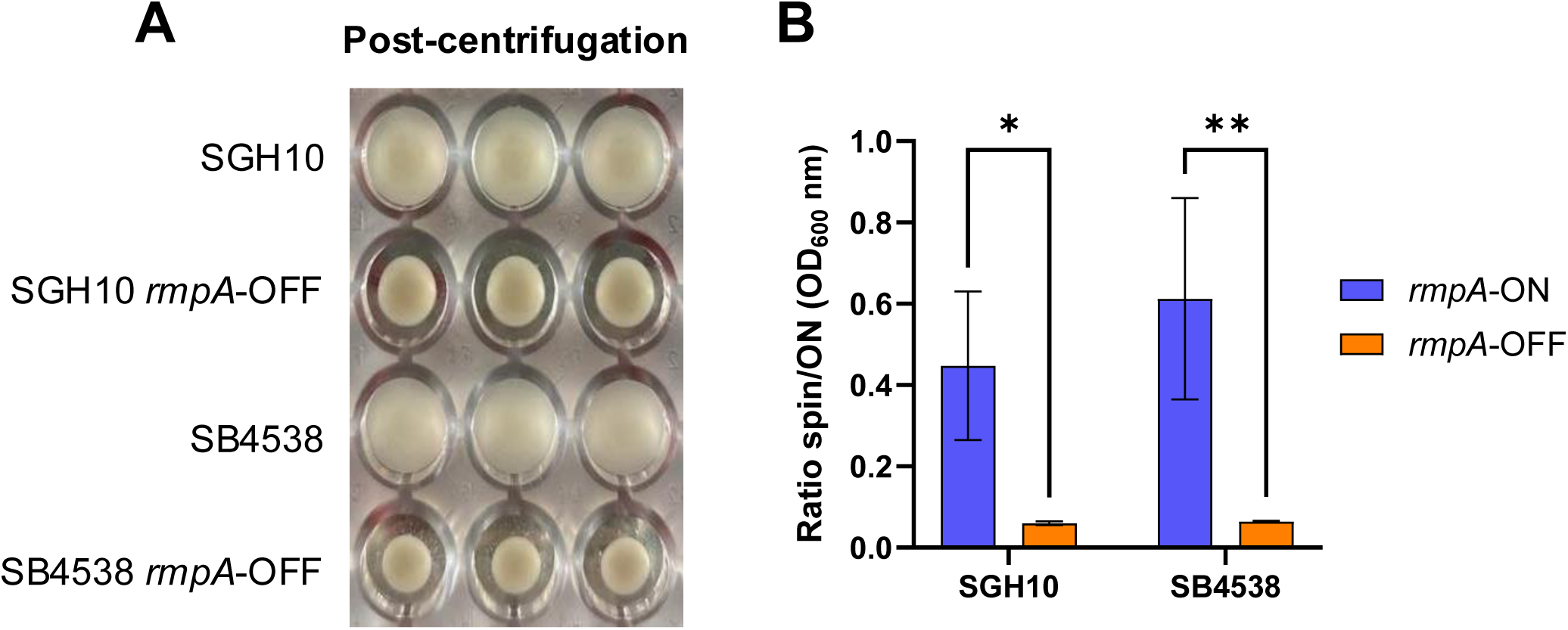
Validation of a high throughput sedimentation assay to measure mucoidy in 96-well microtiter plates. (A) Image of the base of a 96 well plate inoculated with WT SGH10 and SB4538 and their isogenic *rmpA*-OFF mutants. (B) Quantification of sedimentation resistance by measuring the ratio of culture supernatants before and after low-speed centrifugation. Data are expressed as the means and standard deviation of 3 independent experiments. Statistical significances were determined by one-way ANOVA. **; P>0.005, *; P>0.05.

**Figure S2.**
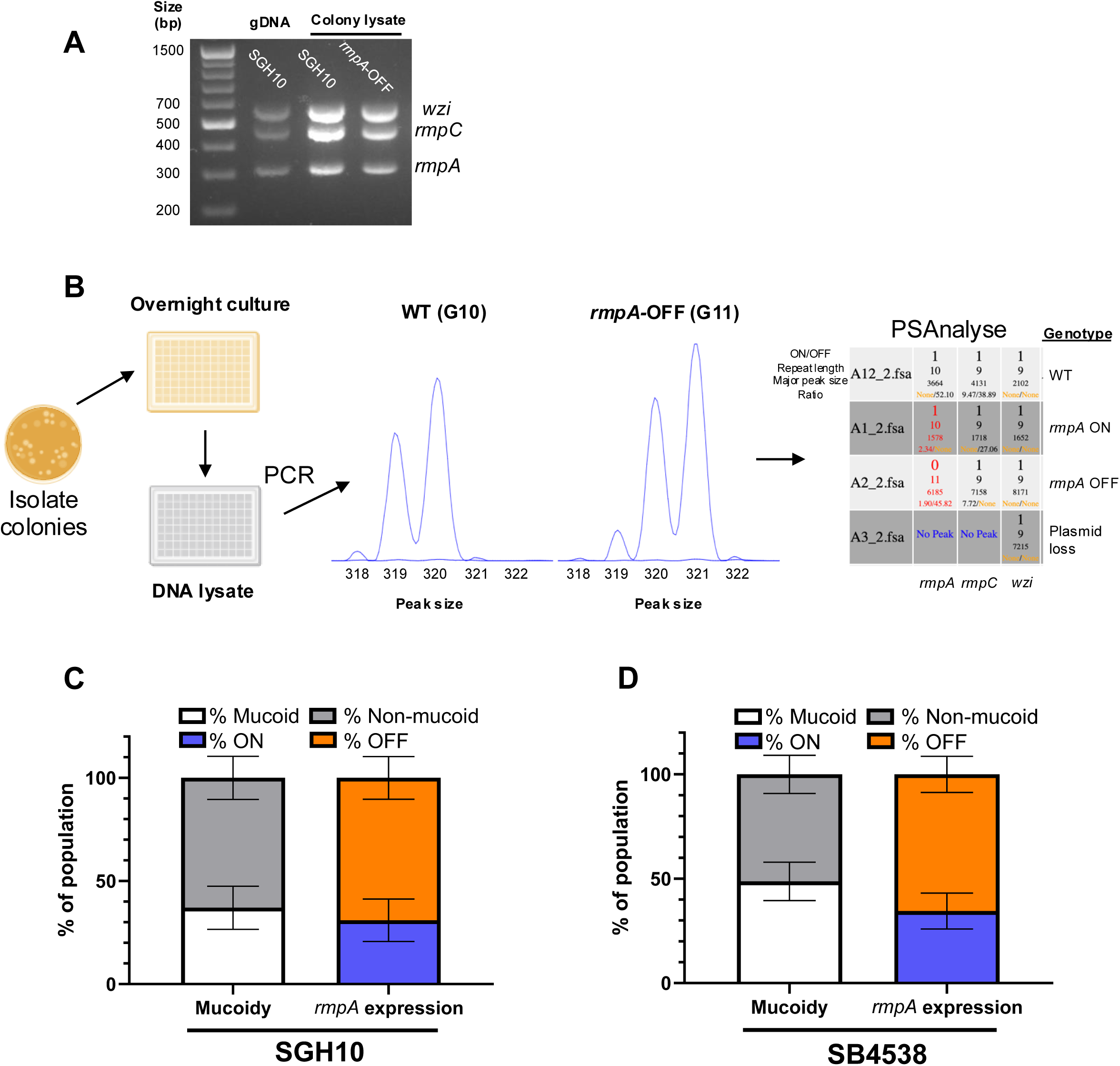
A PCR fragment size assay to measure *rmpA* phase variation. (A) Representative 1% agarose gel showing multiplex amplification of *rmpA*, *rmpC*, and *wzi* genes from strain SGH10 using purified DNA and colony lysates. (B) Experimental workflow to genetically and phenotypically measure the frequency of *rmpA* phase variation in multiple colonies by parallel low speed centrifugation, generation of DNA lysates, multiplex fragment size analysis of repeat-containing regions and automated calling of repeat lengths and ON/OFF expression states with PSanalyse. (C-D) Comparison of the percentage of the population of bacteria called as mucoid (white bars), and non-mucoid (grey bars) by low speed centrifugation with the percentage called as *rmpA*-ON (orange) and *rmpA*-OFF (blue) using the fragment size analysis for SGH10 (C) and SB4538 (D) following combining a 50:50 mix of WT and *rmpA*-OFF mutant overnight cultures derived from single colonies.

**Figure S3.**
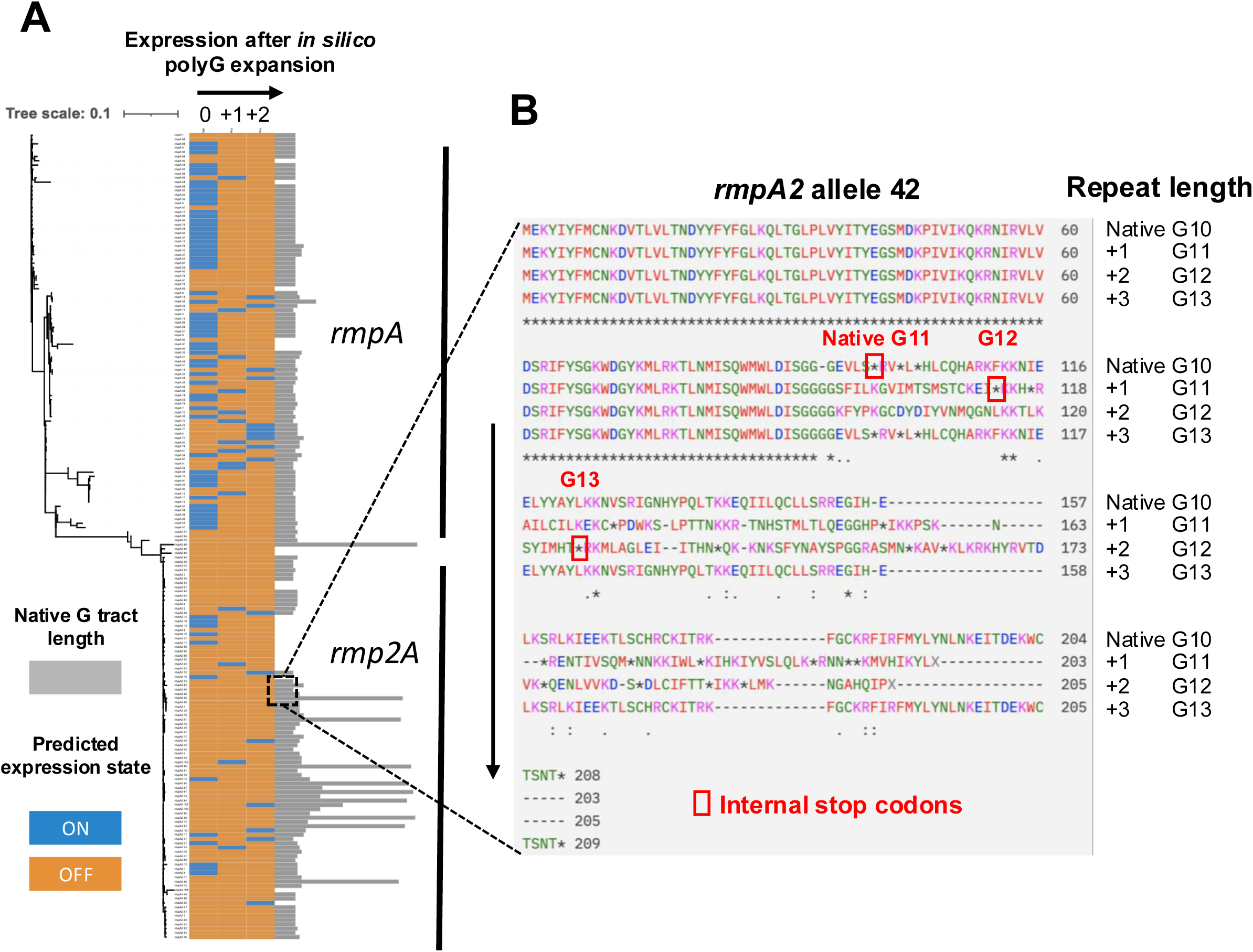
Poly-G repeat polymorphism and truncation in *rmpA2*. (A) Maximum likelihood phylogenetic tree derived from alignment of all available *rmpA* and *rmpA2* alleles from bigsDB (accessed 05/01/2024). ON/OFF expression states are shown by orange (ON) and blue (OFF) bars, following *in silico* expansion of the poly-G repeat as in Figure 6. The length of the poly-G repeat is shown using the grey bars. (B) Alignment of the translated sequences of *rmpA2* allele 42 following *in silico* poly-G expansion. Truncated alleles were translated following *in silico* expansion of the poly-G repeat and predicted amino acid sequences were aligned with clustalOMEGA. The presence of multiple internal stop codons within the coding region after the poly-G repeat irrespective of reading frame are highlighted with red boxes.

**Figure S4.**
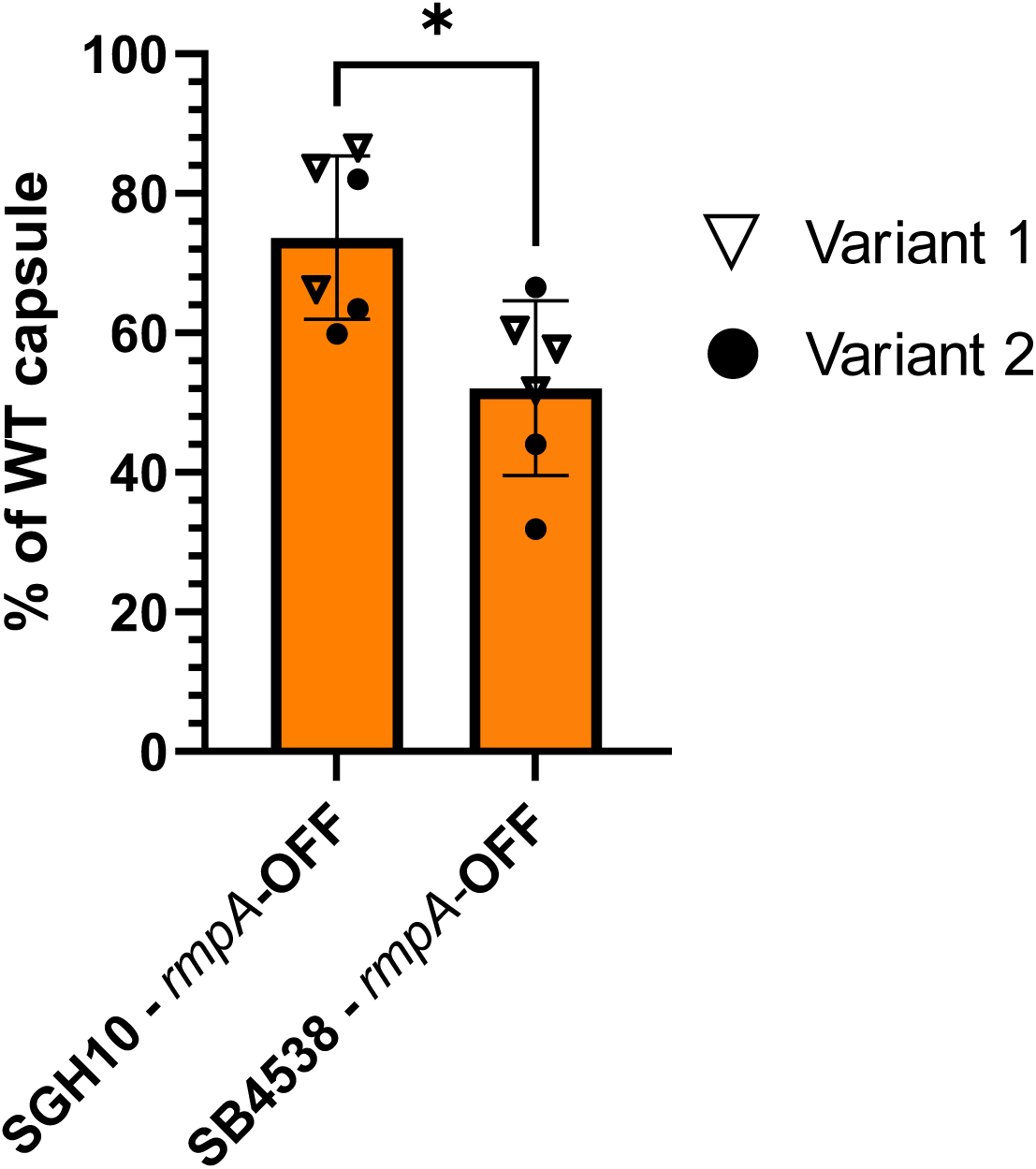
Normalised expression of capsule uronic acid content in *rmpA*-OFF variants compared to their isogenic WT ancestor strains. Capsule preparations were derived from two independent *rmpA-*OFF mutants in both the SB4538 and SGH10 strain backgrounds each in biological triplicate. The relative concentration of uronic acid content in each culture is shown normalised to their respective WT strain. Statistical significance was determined using a student’s T test. *; P>0.05.

**Figure S5.**
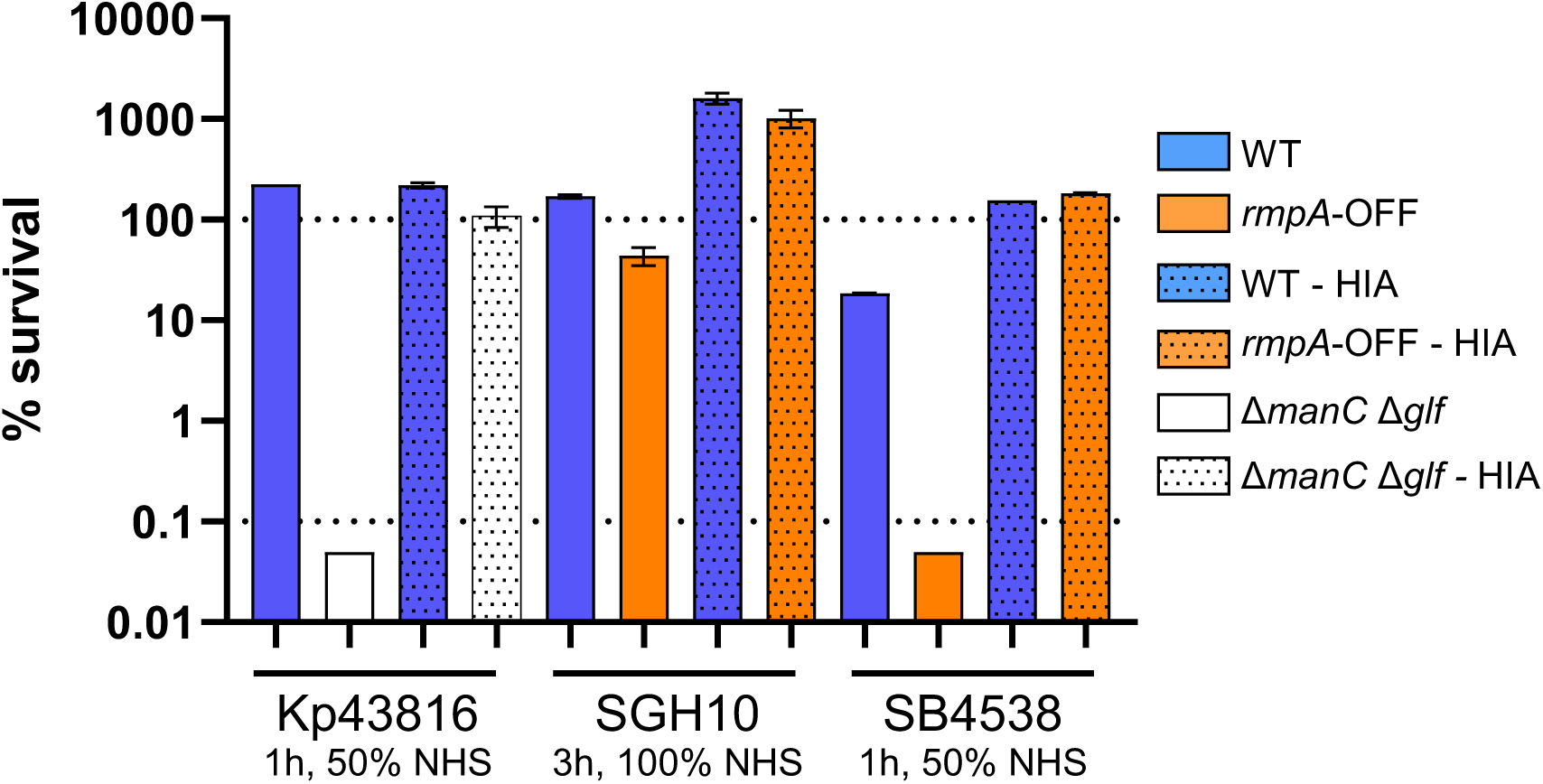
Susceptibility of WT strains, *rmpA-OFF* mutants, and controls strains to human serum with and without heat inactivation. WT strains are shown by blue bars, whereas *rmpA-*OFF mutants are shown by orange bars. A capsule and O-antigen double mutant (Δ*glf*Δ*manC*) o strain Kp43816 was used as a serum sensitive control strain (white bars). Heat inactivated serum samples were used as negative controls for complement activity and are indicated by dotted bars.

**Figure S6.**
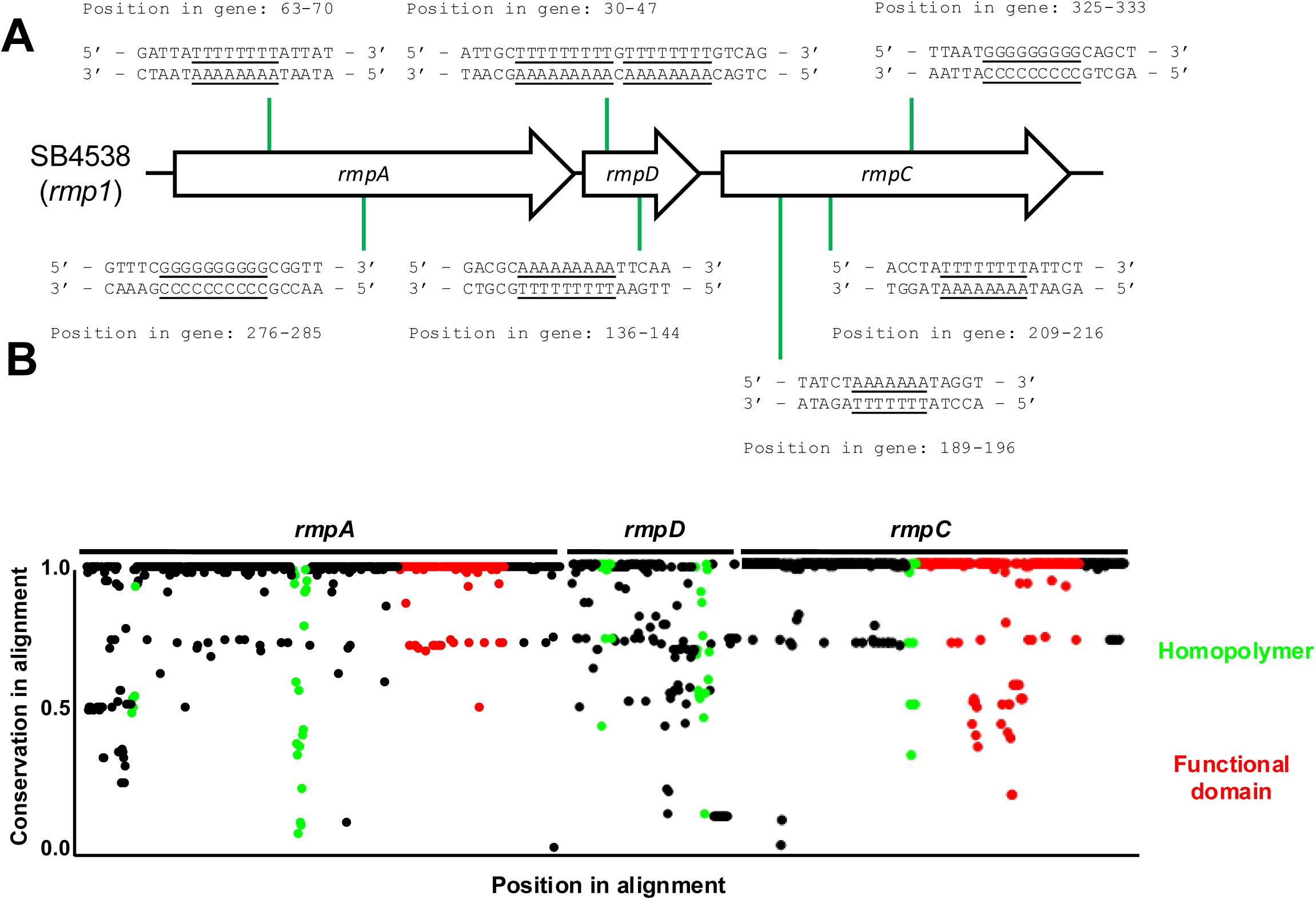
Conservation of homopolymeric repeats in the *rmp* locus. (A) Schematic depicting the *rmp* locus from strain SB4538 annotated with the location and sequence of homopolymeric repeat tracts. (B) Conservation map and repeat tract polymorphism of the RmST typing scheme and location of repeats and functional domains. Regions of sequences containing homopolymers are colored green, whereas regions encoding predicted functional domains as defined by BLASTp analysis of translated amino acid sequences are colored in red.

**Figure S7.**
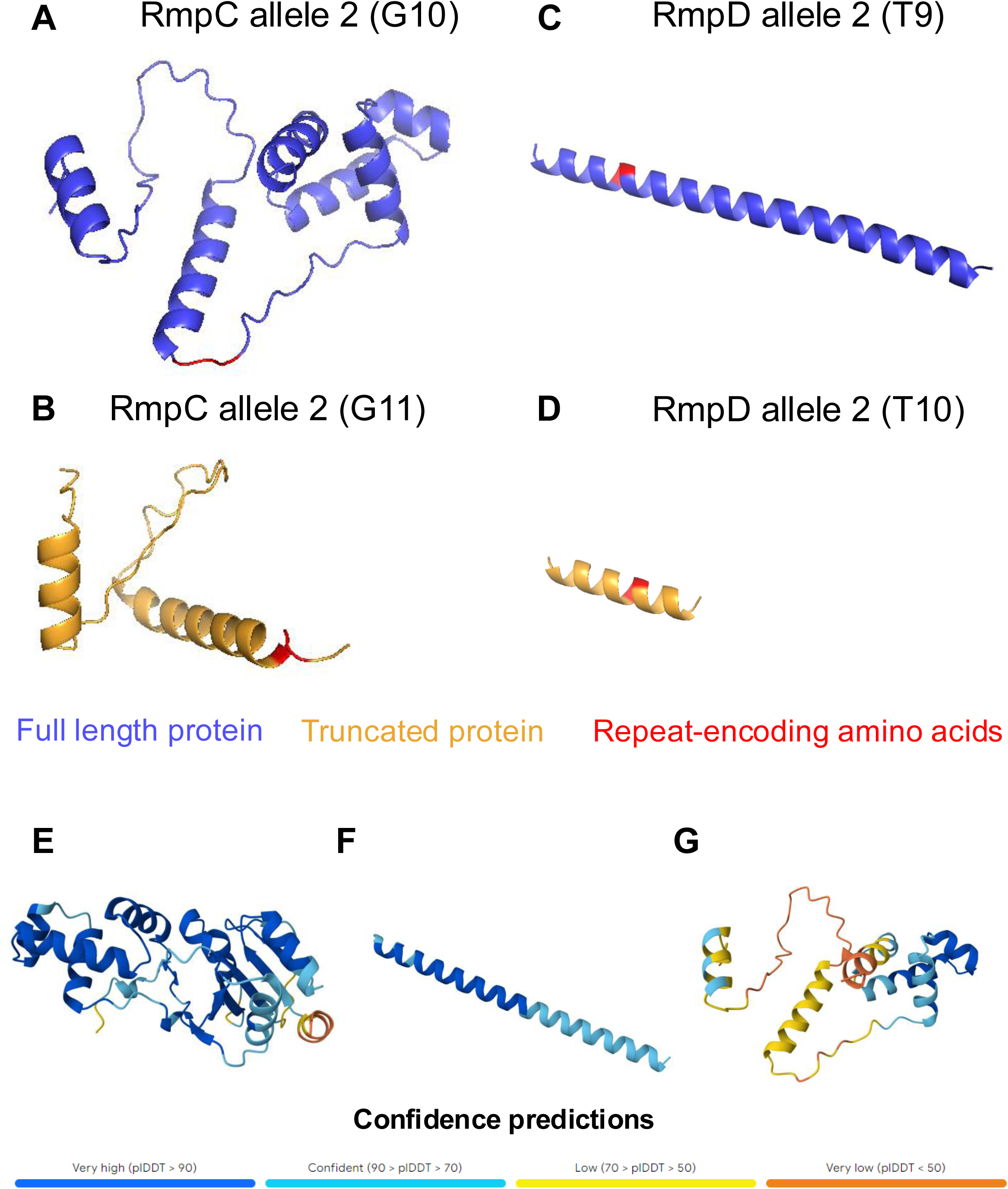
Alphafold structures of WT RmpC, RmpD, and repeat-mediated truncation variants. (A) Structure of RmpC encoded by *rmpC* allele 2. (B) Structure of a RmpC allele 2 G11 truncation variant. (C) Structure of RmpD encoded by *rmpD* allele 2. (D) Structure of a RmpD allele 2 T10 truncation variant. (E-G) Confidence predictions for alphafold structures.

